# A collaboration between immune cells and the choroid plexus epithelium in brain inflammation

**DOI:** 10.1101/2023.08.07.552298

**Authors:** Huixin Xu, Peter Lotfy, Sivan Gelb, Aja Pragana, Christine Hehnly, Frederick B. Shipley, Miriam E. Zawadzki, Jin Cui, Liwen Deng, Milo Taylor, Mya Webb, Hart G. W. Lidov, Mark L. Andermann, Isaac M. Chiu, Jose Ordovas-Montanes, Maria K. Lehtinen

**Affiliations:** Department of Pathology, Boston Children’s Hospital and Harvard Medical School, Boston, MA 02115, USA; Division of Gastroenterology, Hepatology, and Nutrition, Boston Children’s Hospital, Boston, MA 02115, USA; Graduate Program in Biological and Biomedical Sciences, Harvard Medical School, Boston, MA 02115, USA; Broad Institute of MIT and Harvard, Cambridge, MA 02142, USA; Graduate Program in Biophysics, Harvard University, Cambridge, MA 02138, USA; Harvard MD-PhD Program, Harvard Medical School, Boston, MA 02115, USA; Department of Immunology, Blavatnik Institute, Harvard Medical School, Boston, MA 02115, USA; Harvard College, Harvard University, Cambridge, MA 02138, USA; Division of Endocrinology, Diabetes, and Metabolism, Department of Medicine, Beth Israel Deaconess Medical Center, Boston, MA 02215, USA; Department of Neurobiology, Harvard Medical School, Boston, MA 02115, USA; Harvard Stem Cell Institute, Cambridge, MA 02138, USA; Ragon Institute of MGH, MIT and Harvard, Cambridge, MA 02139, USA

## Abstract

The choroid plexus (ChP) is a vital brain barrier and source of cerebrospinal fluid (CSF). Here, we use chronic two-photon imaging in awake mice and single-cell transcriptomics to demonstrate that in addition to these roles, the ChP is a complex immune organ that regulates brain inflammation. In a mouse meningitis model, neutrophils and monocytes accumulated in ChP stroma and surged across the epithelial barrier into the CSF. Bi-directional recruitment of monocytes from the periphery and, unexpectedly, macrophages from the CSF to the ChP helped eliminate neutrophils and repair the barrier. Transcriptomic analyses detailed the molecular steps accompanying this process, including the discovery of epithelial cells that transiently specialized to nurture immune cells, coordinate their recruitment, survival, and differentiation, and ultimately, control the opening/closing of the ChP brain barrier. Collectively, we provide a new conceptual understanding and comprehensive roadmap of neuroinflammation at the ChP brain barrier.

## INTRODUCTION

Brain barriers including the blood-brain barrier (BBB) and the blood-cerebrospinal fluid (CSF) barrier protect the central nervous system (CNS) from systemic insults [1–3]. In conditions of acute inflammation, blood-borne pathogens including viruses (e.g., HIV, Zika, SARS-Cov-2) [4–8], bacteria (e.g., gram-positive and -negative bacteria) [9–11], and parasites (e.g., *Trypanosoma brucei*) [11–13] can directly gain access to the CNS via the brain borders [14, 15]. Subsequent inflammation from infection is associated with an exponentially growing list of lifelong neurologic conditions ranging from microcephaly due to Zika virus exposure [16] and hydrocephalus due to meningitis [17], to persisting cognitive impairments associated with long-COVID [18]. Persisting CNS inflammation is also common to chronic, otherwise sterile conditions such as Amyotrophic lateral sclerosis [19] and Alzheimer’s disease [20]. Elevated immune cell counts in the CSF are well-documented for many neurologic conditions [21–25]. As such, there is a pressing need to increase our understanding of how the brain responds to and resolves inflammation.

The choroid plexus (ChP), located within each brain ventricle and commonly linked to CSF secretion, represents the principal blood-CSF barrier and can provide immediate, far-reaching access throughout the CNS [14, 26]. The human ChP is reported to have a surface area of approximately 50% of the BBB [27]. Sheets of ChP epithelial cells enclose a vascularized stroma consisting largely of mesenchymal and resident immune cells [28]. Resident macrophages (epiplexus / Kolmer cells) are also located on the apical, CSF-contacting surface of the ChP, where they presumably patrol the ChP and the brain’s ventricles [15]. Recent studies are revealing functions for ChP cells ranging from fluid and protein secretion to gating the entry of peripheral signals including immune cells into the CNS during inflammation [1, 2, 25, 29, 30]. Recent transcriptome and fate-mapping analyses uncovered a transient expansion of resident and recruited macrophages at the brain borders including the ChP following *Trypanosoma brucei* exposure [12]. Curiously, inflammation of the ChP itself is also emerging as a common feature of seemingly unrelated neurologic conditions as diverse as hydrocephalus [31], schizophrenia [32], Amyotrophic lateral sclerosis [33], and Alzheimer’s disease [34]. Pro-inflammatory transcriptional signatures in ChP from COVID-19 patients have been linked to chronic CNS inflammatory conditions [35]. Despite these observations, the mechanisms by which the ChP brain barrier coordinates a productive response to protect the brain from either acute or chronic inflammation is not well understood.

Clues to understanding how the ChP addresses inflammation and tissue remodeling can be gleaned from general principles of inflammation established in the body’s other internal organs including visceral organs, lung, and heart. Inflammation in response to infection or sterile injury typically triggers the activation of resident macrophages, which together with other cells, stimulate the endothelium to upregulate adhesion molecules that induce the stepwise recruitment of additional cell types. Neutrophils are often the first to arrive to clear pathogens and debris. Monocytes are subsequently recruited and differentiate into macrophages to promote tissue remodeling and repair [36–38]. A similar series of steps takes place in the brain parenchyma following acute traumatic injury [39]. In the case of meningitis, suppression of macrophage responses by meningeal nociceptors releasing calcitonin gene-related peptide CGRP reduced neutrophil recruitment and worsens the disease [40]. Despite the general understanding that the ChP barrier can recruit peripheral immune cells into the brain, progress in understanding the mechanisms and steps involved, and the role(s) of the ChP, in this process is limited. One main roadblock is the lack of tools available for visualizing and manipulating the ChP barrier *in vivo*.

Here, we advance our understanding of ChP regulation of inflammation by leveraging recent breakthroughs in chronic two-photon imaging of ChP in awake mice [41] and single cell transcriptomics [42] in a model of meningitis. Bacterial meningitis can result from invasion into the brain of any of several species of bacteria (e.g., *N. meningitidis, S. pneumoniae, S. agalacie*) and is clinically diagnosed in part by high leukocyte / white blood cell count in the CSF [43]. We modeled meningitis in mice by intracerebroventricular (ICV) delivery of lipopolysaccharides (LPS), a bacterial endotoxin found in essentially all Gram-negative bacteria. Its bioactive component, lipid A, leads to the activation of the innate immune system via Toll-like receptor 4 (TLR4) signaling, making it a good tool for modeling meningitis in mouse [44]. Thus, we used chronic two-photon imaging to track the real-time ChP recruitment of leukocytes and monocytes following ICV of LPS. Live imaging uncovered an entirely new, bidirectional recruitment of immune cells to the ChP, including macrophages to the ChP from the CSF in addition to monocytes and neutrophils from the periphery. This unique recruitment pattern boosted the available resident macrophages, especially those at the apical surface (the epiplexus cells) and promoted resolution of inflammation and healing of epithelial barrier. Informed by imaging, single cell transcriptome analyses of CSF and ChP performed at key steps of the process revealed a molecular map and stepwise mechanisms for immune cell recruitment, survival, differentiation, and barrier repair. Ultimately, a specialized and transient population of epithelial cells led the way, coordinating immune cell recruitment (by upregulating key chemokines), nourishing immune cells and favoring their differentiation into macrophages (transient upregulation and secretion of colony-stimulating-factor CSF1), and opening/closing of the ChP barrier (orchestrated matrix metalloprotease expression, upregulation of adhesion molecules for securing tissue-repair macrophages). Collectively, our data demonstrate that there is a collaborative and synergetic relationship between immune cells and the ChP epithelial cells. The ChP functions as an immune organ that actively addresses and repairs the brain barrier during inflammation akin to other internal body organs. Our work provides a comprehensive roadmap and a new conceptual understanding of neuroinflammation at the ChP brain barrier.

## RESULTS

### Choroid plexus is a central hub of brain immune cell activity in bacterial meningitis

Examination of ChP samples from patients with bacterial meningitis (18-year-old female, 10-day-old male, and 2-week-old male) revealed a striking number of CD45 and CD163 immunoreactive cells compared to controls (**Figure 1A, Figure S1A**). Indeed, the ChP was swollen and contained numerous peripheral leukocytes of high CD45-positivity that appeared to extravasate into the brain ventricles (**Figure 1B, Figure S1B**). The inflamed ChP tissues also harbored increased numbers of amoeboid, activated CD163+ macrophages (**Figure 1C, Figure S1C**), altogether indicating robust immune activation at the ChP-CSF barrier during meningitis.

**Figure 1.**
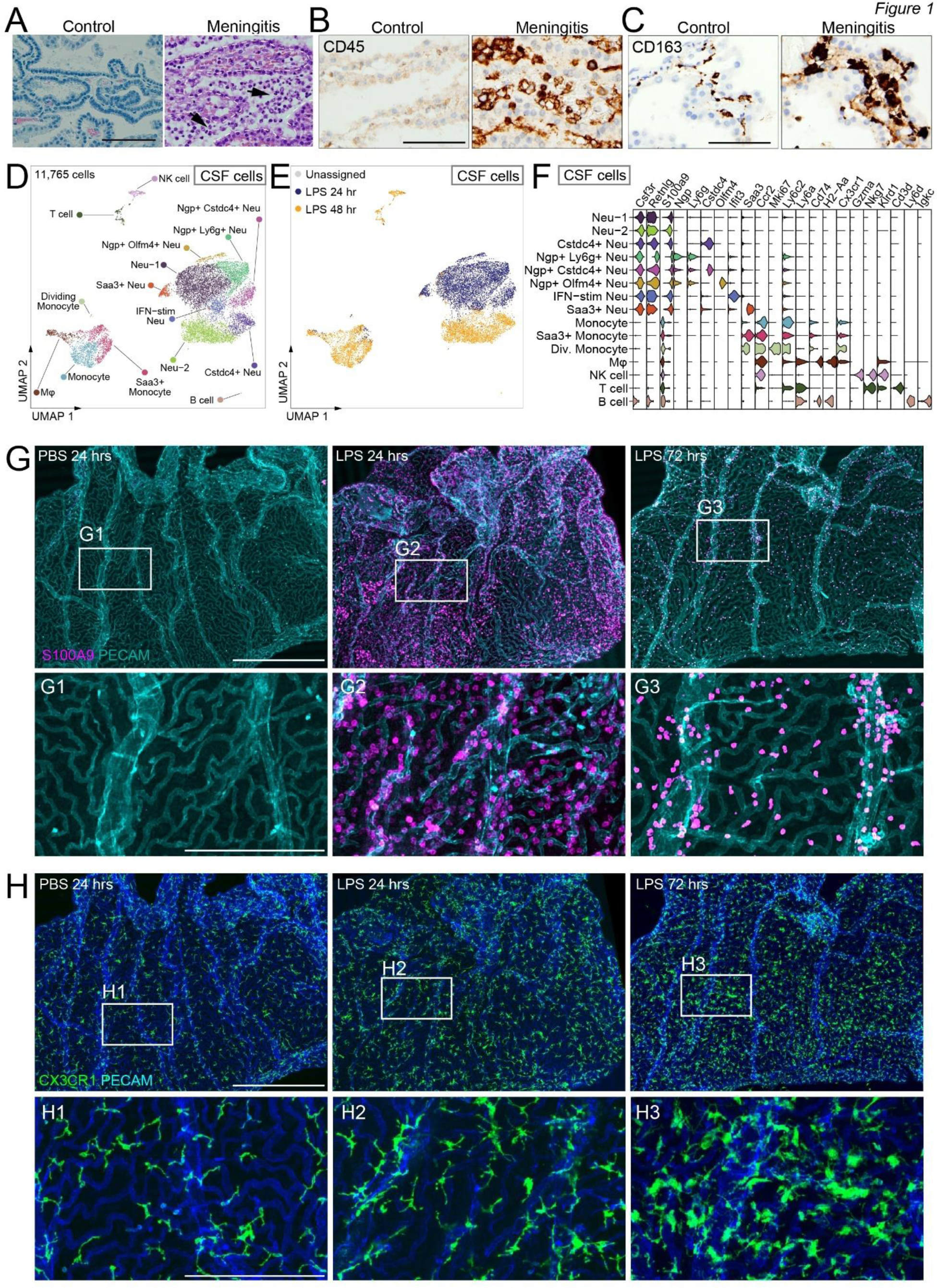
The ChP is a key site of inflammation in meningitis. **(A-C)** Pathology specimens from patients with meningitis vs. controls analyzed by H&E (**A**, with ChP is outlined in the meningitis case and black arrows denote CSF leukocytes), CD45 staining for infiltrated immune cells (**B**), and CD163 staining for macrophages (**C**). Scale = 100 µm. (**D-H**) Analyses of mouse model of meningitis. (**D**) UMAP embedding of 11,765 immune cells collected from CSF 24 hours or 48 hours following intracerebroventricular injection (ICV) LPS challenge for scRNAseq. Neu = Neutrophil; MΦ = Macrophage; IFN-stim = Interferon-stimulated. (**E**) UMAP colored by hash call assignment to each treatment group with pooled samples from 5 adult mice, including those without definitive hash identities (”Unassigned”). (**F**) Violin plot of representative genes used to assign cell type and state identity to clusters. (**G**) Representative images showing accumulation of S100A9+ leukocytes by wholemount ChP explant histology. Scale = 500 µm (200 µm in G1-G3). **(H)** Representative images from wholemount explant histology, showing increasing amount of CX3CR1+ macrophages with amoeboid morphology following LPS. Scale = 500 µm (200 µm in H1-H3).

To elucidate the path that these cells take to enter the brain and related regulatory processes in the ChP, we modeled acute brain infection-associated inflammation in mice by a single intracerebroventricular (ICV) injection of lipopolysaccharides (LPS, *E. coli*), a major component of the cell walls of gram-negative bacteria. In human patients of bacterial meningitis, influx of leukocytes in the CSF is a key diagnostic standard [45]. Using single-cell RNA sequencing (scRNAseq) we confirmed robust cellular infiltration into the CSF 24 hours following LPS delivery (**Figure 1D-F, Figure S2A-D, Supplemental Dataset 1**). Leukocytes in mouse CSF 24 hours following LPS were predominantly neutrophils (*Csf3r*, *Retnlg*). These cells formed six distinct clusters including three *Ngp*+, *Ly6g*+ clusters, one *Saa3+* cluster, and one *IFN*-stimulated cluster, consistent with reported diverse neutrophil populations following infection [46, 47]. By 48 hours, myeloid cells including monocytes and macrophages (*Ccr2, Ly6c2, Cd74, H2-Aa*) became the predominant CSF cell type. We confirmed the accumulation of these major cell classes at the ChP by immunostaining (**Figure 1G-H**). Increased S100A9+ leukocytes, CX3CR1+ macrophages, and CD45-high leukocytes were recapitulated with heat-killed and live *Streptococcus agalactiae* (Group B *Streptococcus,* GBS), a gram-positive bacterial pathogen that causes meningitis [40], delivered ICV (**Figure S3A-B**). These data suggest the immune activation is not unique to LPS but likely a general response to multiple common bacterial infections. Notably, whole brain sections revealed many Ly6G+ neutrophils in the ChP, with only small numbers of neutrophils observed within the subventricular zone and the septum (**Figure S3C**, arrows), suggesting the ChP is the primary site of immune cell infiltration into the brain in this model. More Ly6G+ cells were observed at the “base” of the ChP connecting with the brain parenchyma than the top free margin in the CSF, suggesting the cells may enter the ChP from the base and migrate towards the top.

### Choroid plexus leverages temporally and spatially distinct immune activation programs

To profile ChP immune activation temporally and spatially, we leveraged longitudinal two-photon *in vivo* imaging and visualized real-time leukocyte accumulation at the ChP and subsequent leukocyte entry into the brain (**Figure 2A**). To visualize peripheral leukocytes, mice harboring the ZsGreen reporter (JAX Ai6) were crossed with mice expressing Cre recombinase under the control of the lysozyme 2 (Lyz2) promoter (B6.Cg-Gt(ROSA)26Sortm6(CAG-ZsGreen1)Hze/J) to generate Lyz2-ZsGreen progenies. While resident ChP macrophages express Lyz2, our whole-mount explant histology confirmed the vast majority of Lyz2-ZsGreen cells in ChP 24 hours following LPS co-expressed S100A9 (59.28% ± 13.65%, N=3; **Figure S4A**), a marker for infiltrating neutrophils and monocytes. Longitudinal imaging of the same ChP subregion before and after LPS exposure captured the full spectrum of leukocyte infiltration at the ChP from a very low level at 8-12 hours following LPS, when occasional cells appeared in the ChP, to a “flood” of infiltrated leukocytes by 24-28 hours, in the ChP and CSF (**Figure 2B-D, Supplemental videos 1 and 2**). We also identified discrete “hot spots” of the ChP-CSF interface where Lyz2+ leukocytes extravasated into the ChP, then burst across the epithelial barrier, steaming into the CSF (**Supplemental video 2**). These data suggest that immune infiltration may have spatial preferences.

**Figure 2.**
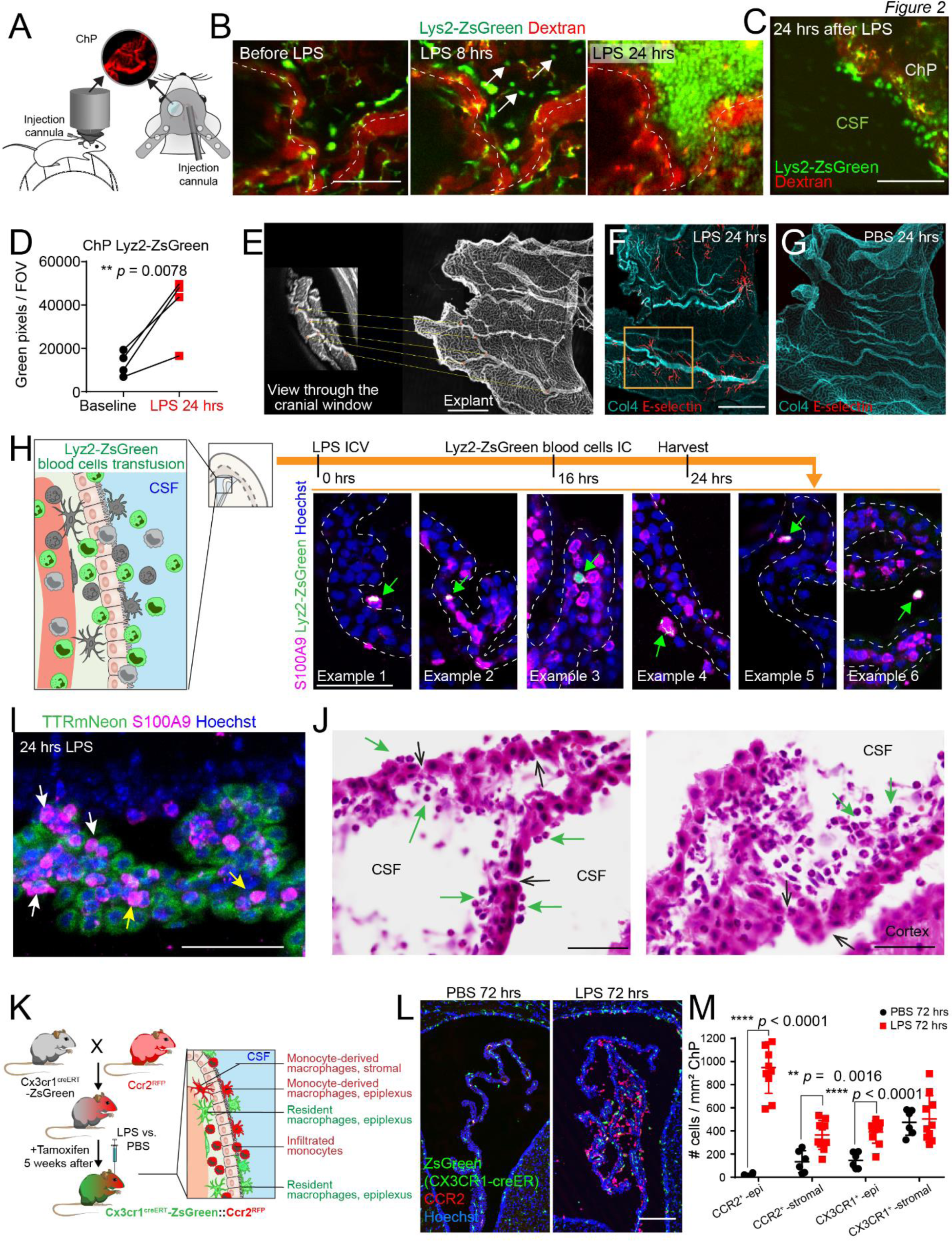
The ChP allows rapid leukocyte infiltration across its epithelial barrier into the CSF. (**A**) Schematic of the cranial window, head post, and injection cannula that allow live two-photon recording of the ChP in awake mice coupled with ICV injection. (**B**) Representative still images extracted from 3 serial *in vivo* imaging sessions of the same mouse, showing the gradual infiltration of Lyz2+ leukocytes into the ChP over the course of 24 hours following LPS ICV delivery. Dotted lines denote major vasculature. See also Supplemental video 1. Scale = 100 µm. (**C**) Representative still images extracted from *in vivo* imaging showing high numbers of Lyz2+ infiltrating leukocytes both in the ChP and in the CSF, 24 hours following LPS delivery. See also Supplemental video 2. Scale = 100 µm. (**D**) Quantification showing significant increase of Lyz2+ signal from baseline to 24 hours following LPS (** *p* = 0.0078, N=3). (**E**) Representative image demonstrates the spatial correspondence between *in vivo* view and wholemount explant. Scale = 500 µm. (**F-G**) Representative image of a wholemount explant showing selective expression of E-selectin. Scale = 500 µm; Yellow square indicates typical *in vivo* field of view. (**H**) Schematics and 6 representative images demonstrating infiltration of Lyz2-ZsGreen S100A9+ immune cells from peripheral blood into the ChP (examples 1-4) and CSF (examples 5-6). Scale = 50 µm. (**I**) Representative image showing S100A9+ peripheral immune cells accumulating within the ChP stroma (yellow arrow) and penetrating epithelial layers (white arrow). Epithelial cells are labeled by endogenous TTR^mNeon^. Scale = 50 µm. (**J**) Representative H&E images from mice 24 hours following LPS ICV delivery showing presence of neutrophils in both ChP stromal space and CSF-facing epiplexus surface (green arrows), as well as breakage in epithelial bi-layers (black arrows). Scale = 50 µm. (**K**) Schematic depicting breeding scheme to acquire mice harboring CCR2^RFP^ and tamoxifen-inducible Cx3cr1^Zsgreen^, which distinguishes between infiltrated monocytes and those that derived into macrophages (RFP+, ZsGreen-) and resident macrophages (ZsGreen+), in both stromal and epiplexus spaces of the ChP. (**L-M**) Representative images and quantification showing significant increase in epiplexus (**** *p* < 0.0001) and stromal (** *p* = 0.0016) RFP+ cells, as well as a small, but significant, increase of epiplexus ZsGreen+ cells (**** *p* < 0.0001). Scale = 100 µm. All quantitative data are presented as mean ± SD.

Following *in vivo* imaging, we prepared whole mounts of the ChP and used its characteristic vasculature to guide *post hoc* analyses of the imaging locations. The two-photon imaging field of view contained the caudal aspect of the anterior domain of the lateral ventricle ChP (**Figure 2E**). We observed robust endothelial expression of E-selectin 12-24 hours following LPS in the ChP, including regions within the two-photon imaging field of view (**Figure 2F-G**). E-selectin expression is consistent with a post-capillary venule localization, likely indicating sites of leukocyte extravasation from the blood into the ChP before bursting through the epithelial barrier.

Further supporting the concept that the ChP is a crucial hub of leukocyte activity following LPS, we found that peripherally labeled leukocytes entered the ChP and were closely positioned on both the basal, stromal side and apical, CSF side of the epithelium. First, we collected peripheral blood leukocytes from LPS-stimulated Lyz2-ZsGreen mice, transfused these cells into the blood of LPS-stimulated WT mice by intracardiac injection, and subsequently detected Lyz2-ZsGreen cells within ChP stroma, on the ChP surface (**Figure 2H, examples 1-4**), and in the CSF (**Figure 2H, examples 5-6**). Similarly, we labeled blood-sourced neutrophils by intracardiac delivery of anti-Ly6G antibody conjugated to Alexa-594, which appeared within the ChP shortly after, as shown by two-photon *in vivo* imaging (**Figure S4B, Supplemental video 3**). Second, we performed histological analyses to visualize peripheral leukocyte infiltration through the epithelial bilayers. *Ttr^mNeon^* mice with fluorescently labeled ChP epithelial cells [48] revealed S100A9+ cells in the ChP stroma, with more in the base than in the top region as described earlier, and also breaching the epithelial layer throughout the ChP 24 hours after LPS ICV (**Figure 2I**). Neutrophils in the ChP stroma and on the apical, ChP-CSF surface were accompanied by breaks in tight junctions between epithelial cells (**Figure 2J**). We also captured the positioning of neutrophils along the blood vessels, in the ChP stroma, and then on the surface of epithelial cells with electron microscopy (**Figure S4C**). Lastly, we applied imaging flow cytometry (ImageStream) to the ChP following LPS ICV to visualize Ly6G/C+ neutrophils/monocytes and epithelial cells (Kiravia dye, “Kirva”) at single-cell resolution after tissue dissociation. Unlike most dissociated cells, many Ly6G/C+ cells remained attached to epithelial cells after tissue digestion (**Figure S4D-E**; 24 hours: 21.6 ± 16.7% of total live cells, N =3, and 48 hours: 5.6 ± 2.1%, N=4), indicating tight physical interactions. Using mice co-expressing *Ccr2^RFP^* and tamoxifen-inducible *CX3CR1^ZsGreen^* which labels only resident macrophages 5 weeks after tamoxifen treatment (**Figure 2K**), we found that infiltrated CCR2+ monocytes accumulated in both stromal and epiplexus spaces following LPS (**Figure 2K-M**). Collectively, these data strongly support the conclusion that peripheral leukocytes infiltrate the brain through the ChP epithelial barrier in response to LPS.

Following the initial wave of infiltrated leukocytes, we tracked the real-time macrophage responses at the ChP 48-72 hours following LPS (**Figure 3A-B, Supplemental video 4**), the period where we observed the most robust macrophage changes by histology (**Figure 1H**). During this time, numbers of visibly motile ChP macrophages were increased drastically, encasing the entire tissue (**Supplemental video 4**). *Cx3cr1^GFP^*;*Ccr2^RFP^* dual labeled mice revealed a significant increase of cells co-labeled with both CX3CR1-GFP and CCR2-RFP at 72 hours following LPS ICV (**Figure 3C-D**), suggesting the infiltrated monocytes differentiated into macrophages and that monocyte-derived macrophages accounted for the majority of newly gained ChP macrophages. Consistent with increased CCR2+ monocytes both in the stromal space and on the epiplexus surface (**Figure 2K-L**) and increased CCR2+/CX3CR1+ macrophages (**Figure 3D**), we also found increased numbers of Iba1+ macrophages throughout the ChP (**Figure 3E-G**).

**Figure 3.**
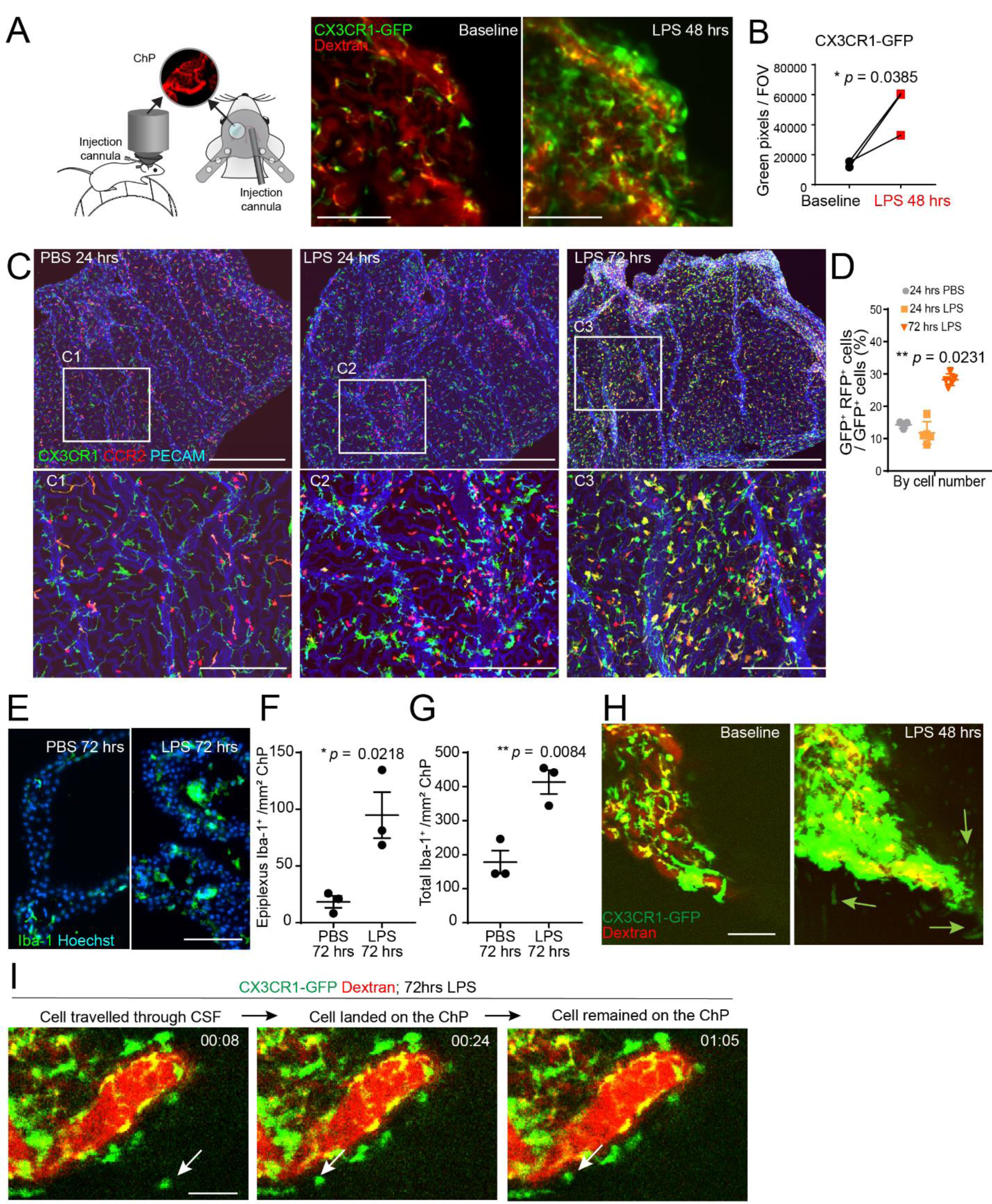
The ChP recruits macrophages from peripheral monocytes and CSF macrophages. (**A**) *In vivo* imaging schematics (same as Figure 2A) and representative still images of *in vivo* imaging showing increasing numbers of ChP macrophages over the course of 48 hours following LPS delivery. Also see Supplemental video 4. Scale = 100 µm. (**B**) Quantification showing significant increase of CX3CR1+ signal over the course of 48 hours following LPS ICV delivery (* *p* = 0.0385, N=3). (**C**) Representative images from wholemount explant histology, showing increasing amount of CCR2+ monocytes and CCR2+/CX3CR1+ macrophages following LPS delivery. Scale = 500 µm (200 µm in C1-C3). (**D**) Quantitative analysis showing increasing number of cells that are CCR2+/CX3CR1+ in ChP wholemount explants. * *p* = 0.0231, one-way ANOVA. (**E-G**) Representative images and quantifications showing increase of epiplexus (F, * *p* = 0.0218) and total (G, ** *p* = 0.0084) Iba1+ macrophages in the ChP following LPS ICV delivery. Scale = 100 µm. (**H**) Representative images compressed from one-hour long *in vivo* imaging, showing increased Cx3cr1+ macrophages traveling through CSF (green arrows) following LPS delivery. Also see Supplemental video 5. Scale = 100 µm. (**I**) Representative images showing CX3CR1+ macrophage traveling from CSF and landing on the ChP. Also see Supplemental video 8. Scale = 50 µm. All quantitative data are presented as mean ± SD.

Two-photon imaging revealed that in addition to blood-sourced monocyte-derived macrophages, epiplexus macrophages arrived at the ChP via the CSF. We recorded substantial numbers of CX3CR1+ cells traveling through CSF following LPS ICV, lasting approximately from 48 to 72 hours following LPS ICV (**Figure 3H, Supplemental video 5**), consistent with results from CSF scRNAseq (**Figure 1D-F**). CSF-macrophages expressed neither microglial signature genes (e.g., *Sall1, P2ry12*) [49] nor barrier-associated macrophage genes (e.g., *Pf4*) [50], suggesting either a peripheral origin or a change of cellular signature upon leaving their native environment in the central nervous system (CNS). In support of the extensive travel of macrophages, we observed dynamic translocation of macrophages from the ventricle wall into the CSF and towards the ventricle wall from the CSF (**Supplemental video 6**), as well as cases of macrophages traveling towards the ChP from CSF (**Supplemental video 7**). Some CSF macrophages landed on the ChP and remained there as epiplexus macrophages (**Figure 3I, Supplemental video 8**).

Collectively, these data establish a time course of ChP immune infiltration and activation following brain LPS exposure that involves both infiltrated leukocytes and dynamic macrophages. We show that peripheral neutrophils and monocytes infiltrated as first responders across spatially distinct regions of the ChP, which allowed the expansion of the ChP macrophage pool from two distinct sources: (1) peripheral monocyte differentiation and (2) CNS macrophage recruitment from the CSF.

### Epithelial cells govern the logical, stepwise inflammatory response at choroid plexus

To determine the cell types and mechanisms involved in regulating ChP immune infiltration revealed by *in vivo* two-photon imaging, we performed scRNAseq on the ChP from LPS-treated mice (**Figure 4, Figure S5A-C, Supplemental Dataset 2**). We identified 2 clusters of neutrophils in the ChP, *Retnlg*+ and *Rpl*-high, from 24 hours and 72 hours post-LPS respectively with representing genes along the spectrum of neutrophil maturity [46]. *Retnlg*+ neutrophils exhibited higher levels of granule genes (*Chil3*, *Lcn2*, *Wfdc21*, *Ngp*, *Mmp9*), inflammatory chemokines (*Ccl3*, *Ccl4*, *Cxcl2*), cytokines (*Il1b*), and mediators (*Nfkbia, S100a8, S100a9*) (**Figure 4A-C**), whereas Rpl-high neutrophils had slightly elevated expression of ribosomal genes (*Rpl18a*, *Rps3*, *Rpl8*) and a signature defined by unique expression of *Siglecf* and *Ptma*, which have been previously described as markers of late-stage neutrophils [51]. The number of neutrophils present at 24 hours exceeded those observed at 72 hours (**Figure S5D**), corroborating findings from CSF scRNAseq (**Figure 1D-F**) and imaging studies. Additionally, we observed a population of *Ccr2+ Plac8+* monocytes infiltrating into the ChP at 24 hours and *Ccr2+ Cx3cr1+* infiltrating macrophages and monocyte-derived dendritic cell (moDC) populations at 72 hours, including cells that are proliferating (Mki67+) (**Figure 4A-C, Figure S5D**). cDC1s were present in the ChP at baseline in concordance with previous reports [28], suggesting that these cells naturally patrol the tissue and may cross-present antigens to promote CD8+ T-cell responses. Accordingly, we observed significant infiltration of CD8+ T cells at 72 hours post-ICV LPS (**Figure 4A-C, E, Figure S5D**).

**Figure 4.**
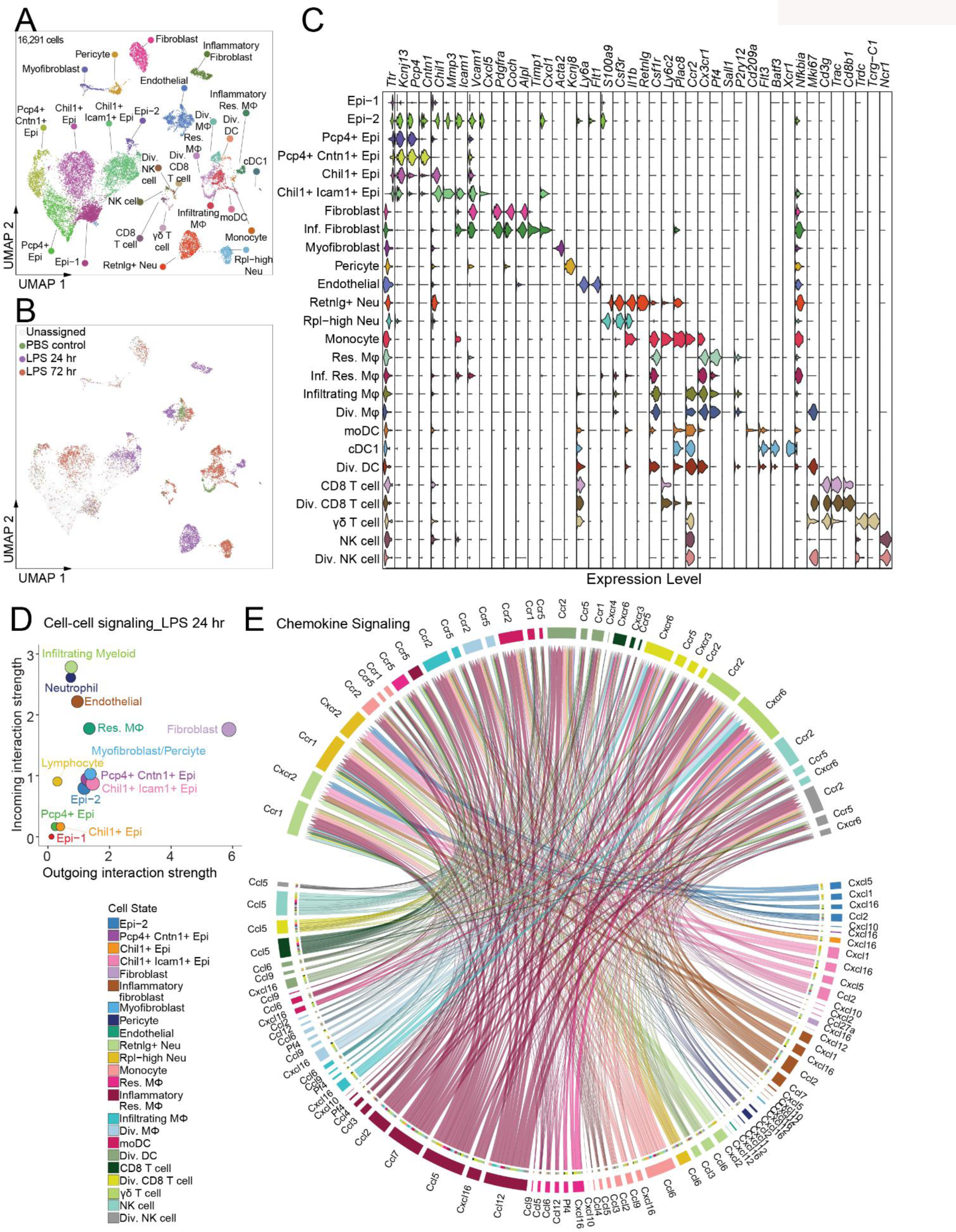
scRNAseq of the inflamed ChP reveals complex and dynamic immune signatures to support immune infiltration. (**A**) UMAP embedding of 16,291 cells collected from ChP 24 hrs or 72 hrs following LPS ICV or 24 hrs following PBS ICV for scRNAseq. Neu = Neutrophil; MΦ = Macrophage; Epi = epithelial cells. (**B**) UMAP as in (A) colored by hash call assignment, including those without definitive hash identities (”Unassigned”). (**C**) Violin plot of differentially expressed marker genes used to assign cell type and state identity to clusters. (**D**) CellChat analysis showing cell-to-cell interactions mediated by secreted ligand (outgoing) and receptor (incoming) pairs at its peak strength 24hrs after LPS ICV. (**E**) CellChat analysis showing chemokine signaling from all cell types (outgoing arrows) to immune cells (incoming arrows).

We observed heightened cell-cell interactions through secretory signaling at 24 hours following LPS (**Figure 4D**), including extensive chemokine signaling from multiple sources within the ChP (**Figure 4E**). We first identified a population of resident macrophages (*Cx3cr1+ Pf4+ Ccr2-*) that displayed an intensely inflammatory transcriptional response including high levels of chemokine expression at 24 hours (*Nfkbia, Ccl2, Ccl5, Ccl7*, **Figure 4A-C, E, Figure S5D**) and reverted back to baseline at 72 hours, indicating their roles in initiating leukocyte recruitment. In addition, we identified one *Chil1*+ *Icam1*+ epithelial state out of 6 clusters of *Ttr+* epithelial cells that was induced at 24 hours and produced high levels of chemokines (*Ccl2, Cxcl5, Cxcl16, Cxcl1, Cxcl10*, **Figure 4E, Supplemental Dataset 2**). One out of two clusters of fibroblasts (*Pdgfra*, *Coch*, *Alpl*, *Col1a1*) was enriched at 24 hours and marked by relatively elevated expression levels of chemokines (*Cxcl1*, *Cxcl16*, *Ccl2*, *Ccl7*, **Figure 4A-C, E, Supplemental Dataset 2**) and acute phase proteins (*Saa1*, *Saa3*) (**Supplemental Dataset 2**). Consistently, we detected high levels of cytokines and chemokines within the CSF from mice 24 hours after LPS injection, and less at 72 hours (non-detectable in mice receiving PBS) (**Figure S6**). In addition to chemokines, the ChP also displayed NFkB signaling in *Chil1*+ *Icam1*+ epithelial cells and IFN signaling in *Chil1*+ *Icam1*+ epithelial cells (*Cxcl10*, *Ifit3*, *Isg15,* **Supplemental Dataset 2**) and endothelial cells associated with LPS 24 hours (*Ifitm3*, *Irf7*, and *Isg15,* **Figure S5E**, **Supplemental Dataset 2**). These cellular changes are consistent with the hypothesis that the ChP responds directly to the presence of pathogenic triggers, such as LPS, within the CSF through its epithelial cells and resident macrophages and engages its endothelial cells and fibroblasts. Indeed, removing all resident macrophages with PLX-5622 (CSF1R inhibitor) prevented LPS-mediated immune infiltration into the ChP (**Figure S7A**). Injections of an equal dose of LPS (2.5 µg) directly into peripheral blood also failed to induce substantial leukocyte infiltration compared to ICV injection (**Figure S7B**).

We identified a unique cluster of inflammatory epithelial cells (*Chil1*+ *Icam1*+ epithelial cells, here on referred to as inf-Epi) that emerged 24 hours following LPS ICV and executed a multitude of interactions with immune cells to coordinate the stepwise progression of ChP inflammation, in addition to secreting chemokines. GO analysis revealed that, in comparison to *Pcp4+ Cntn1+* epithelial cells that were primarily associated with baseline condition, the inf-Epi shifted their functional priority from ion transport and transmembrane transport, which are the most common functions of ChP epithelial cells, towards immune recruitment and support (**Figure S5F-G**). First, inf-Epi, as well as inflammatory fibroblasts, expressed a high level of extracellular matrix remodeling factors (*Mmp3, Timp1*, *Adamts4*) complementing the ones expressed by immune cells (*Mmp8, Mmp9, Mmp14*) (**Figure 5A**). Concurrently, epithelial tight junctions were weakened 24 hours following LPS ICV, as shown by loss of Occludin junctions and disrupted organization of Claudin-2 and ZO-1 (**Figure 5B**), enabling infiltration of leukocytes from ChP to CSF. Next, we found robust colony-stimulating-factor CSF1 signaling from inf-Epi and some inflamed fibroblasts and endothelial cells towards myeloid cells at 24 hours, which reduced by 72 hours (**Figure 5C**). CSF1 supports macrophage differentiation and survival [52–54]. This induction of CSF1 in the ChP corresponded with the infiltration of CCR2+ monocytes (24 hours) and preceded their differentiation to CX3CR1+ CCR2+ macrophages and DCs (72 hours), suggesting an active system at the inflamed ChP to support the survival of infiltrated myeloid cells and guide their differentiation towards macrophages. We confirmed this finding with histology showing increased CSF1 expression in the ChP epithelial cells (**Figure 5D**). We further detected increased CSF1 levels in the CSF 24 hours following LPS ICV by immunoblotting, consistent with the secretion of CSF1 by ChP epithelial cells into the CSF (**Figure 5E**). Neutralizing CSF1 by ICV delivery of antibodies 24 hours following LPS ICV reduced epiplexus CX3CR1+/Iba1+ macrophages at 72 hours, consistent with CSF1’s role in supporting macrophage survival (**Figure 5F-G**). In addition, we found that inf-Epi contributed to ChP macrophage gain (as shown in **Figure 3**) by retaining epiplexus macrophages with increased expression of adhesion molecules, including ICAM1 and VCAM1, as shown both by scRNAseq (**Figure 4C**) and histology (**Figure S8A-B**), consistent with prior reports [12, 55]. We performed CSF macrophage transplant studies with neutralizing antibodies against VCAM1 and ICAM1 to confirm their functions (**Figure 5H**). CSF cells were isolated from tamoxifen-induced CX3CR1-ZsGreen mice 48 hours following LPS ICV and delivered ICV into wild-type mice pretreated with LPS ICV for 24 hours. Before cell transfer, the WT mice received anti-VCAM1 and anti-ICAM1 antibodies or dose-matched rat IgG isotype control by ICV. Brains from WT mice were harvested 48 hours after cell transfer to look for epiplexus ZsGreen+ cells on the ChP. Indeed, we found that mice treated with VCAM1/ICAM1 neutralizing antibodies had ∼50% fewer ZsGreen+ cells on their ChP apical surface, as well as ∼50% fewer total Iba1+ epiplexus macrophages (**Figure 5I-J**). Intraventricular injection of antibodies targeting VLA-4α and LFA-1α, which are leukocyte-expressed integrin receptors for VCAM1 and ICAM1 (**Figure S8C**), reduced numbers of epiplexus Iba1+ macrophages to a comparable extent (**Figure S8D-E**). Collectively, our data demonstrate that ChP epithelial cells, and in particular inf-Epi population, carry out multiple layers of support to coordinate immune actions and transport at the ChP.

**Figure 5.**
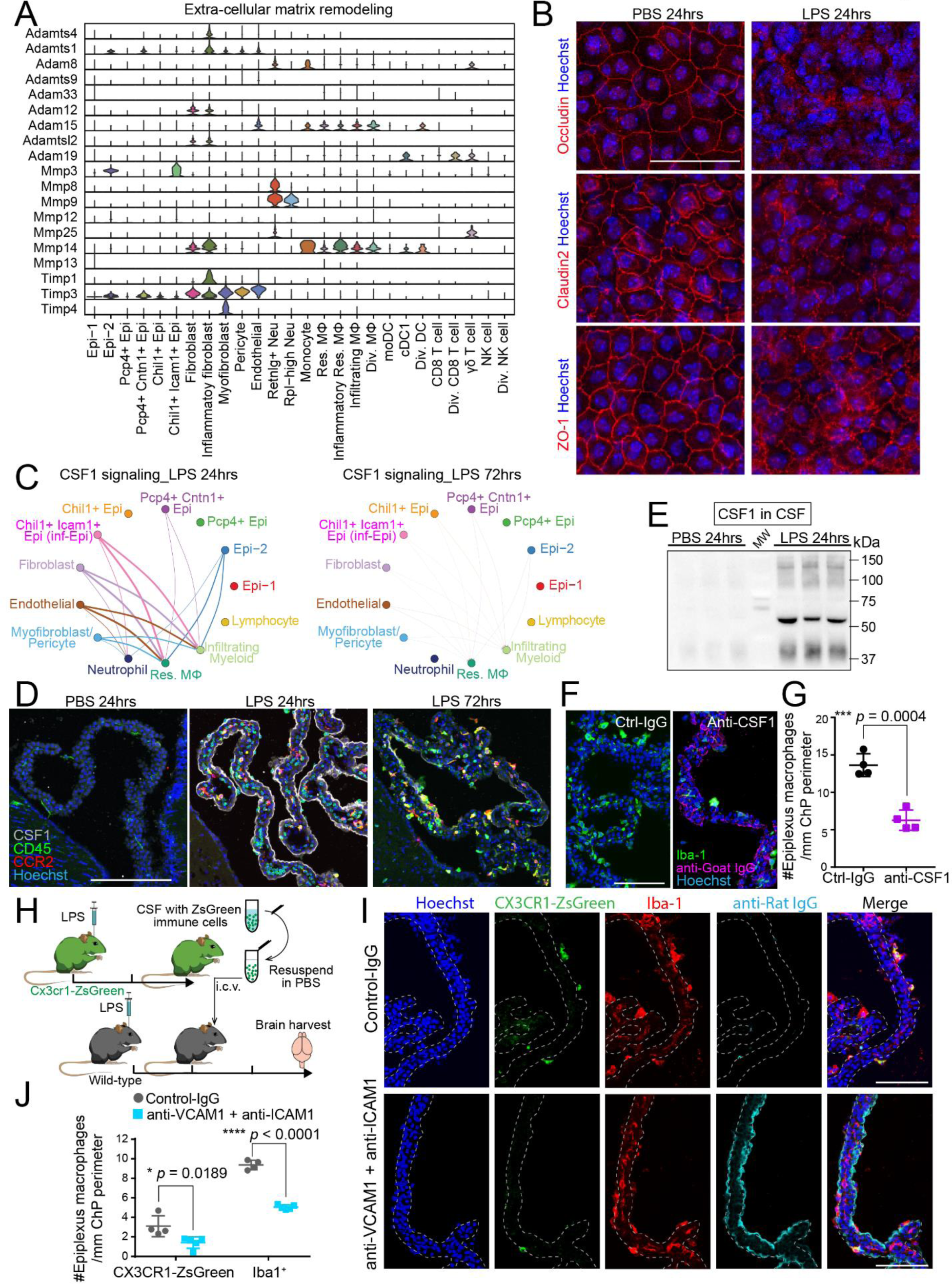
Epithelial cells emerge as key coordinators of choroid plexus immune response. (**A**) Violin plot showing expression of extracellular matrix remodeling proteins by epithelial cells, fibroblasts, and other non-immune cells, as well as neutrophils and macrophages of the ChP. (**B**) Representative images of wholemount ChP explants showing disrupted patterns of occludin, claudin 2, and ZO-1 staining by LPS. Scale = 50 µm. (**C**) Cellchat analysis showing robust CSF1 signaling from ChP epithelial cells, fibroblasts, endothelial cells towards resident and infiltrated macrophages at 24 hours following LPS ICV. (**D**) Representative images showing increased CSF1 expression in the ChP 24 hours following LPS ICV. The expression reduced at 72 hours and was not detectable in PBS ChP; Scale = 200 µm. (**E**) Immunoblot of CSF showing increased level of CSF1 in CSF from mice 24 hours following LPS ICV delivery. 9 µl of CSF was loaded for each lane. (**F-G**) Representative images and quantification showing reduced epiplexus macrophages (Iba1+) in mice treated with anti-CSF1 antibody ICV following LPS ICV. Scale = 100 µm. (**H**) Schematics demonstrating the experimental procedure to collect ZsGreen+ brain resident macrophages within LPS-stimulated donor CSF and transplant to LPS-stimulated recipient mice by ICV. (**I**) Schematics demonstrating how CSF immune cells may be attracted to ChP epithelial cells through VCAM1 - VLA-4α interaction (similar scenarios are expected with ICAM1 – LFA-1α interaction). Blocking either side of the interaction should prevent CSF immune cells landing. (**J-K**) Representative images and quantifications showing reduced numbers of both ZsGreen+ donor CSF macrophages (* *p* = 0.0189) and total Iba1+ epiplexus macrophages (**** *p* < 0.0001) in mice treated ICV with anti-VCAM1/ICAM1 neutralizing antibodies prior to cell transplant. Scale = 100 µm. All quantitative data are presented as mean ± SD.

### Choroid plexus epiplexus macrophages contributed to barrier healing

To determine the functional significance of retaining epiplexus macrophages following LPS-induced inflammation, we first investigated their behaviors using *in vivo* two-photon imaging. We found a range of altered cellular dynamics consistent with phagocytic and pro-healing status. A significant portion of CX3CR1+ macrophages became amoeboid in shape and formed large vacuoles inside their cytoplasm 24 hours after LPS ICV (**Figure 6A-B, Supplemental video 9**). The macrophages also became highly mobile and formed large aggregates throughout the ChP from 24 hours to 72 hours (**Figure 6A-B, Supplemental Video 10**), similar to their reported responses to tissue injury. Further, we specifically observed resident epiplexus macrophages by crossing *Ttr^mNeon^* mice and mice expressing tamoxifen-inducible *Cx3cr1^TdTomato^*. We found that, following LPS ICV, tdTomato+ epiplexus resident macrophages also became highly mobile and motile with large vacuoles in either cell bodies or at the end of processes (**Figure 6C, Supplemental Video 11**), but did not form any large clusters. The differences in observed cellular dynamics suggest both shared and distinguished functions between resident and differentiated macrophages. The number of epiplexus resident macrophages increased over the course of 48 hours (**Figure 6D**), consistent with lineage tracing data (**Figure 2K-M**). To determine the possible content of the vacuoles, we used ImageStream and identified a population of macrophages (F4/80+) that contained Ly6G/C+ neutrophils or monocytes (24 hours: 5.3 ± 2.3% of all live cells, N=3, and 48 hours: 0.6 ± 0.4%, N=4, **Figure S9A-C**), which was further confirmed by immunohistology on brain sections (**Figure 6F**). These findings suggest that some of the macrophages contributed to clearance of neutrophils.

**Figure 6.**
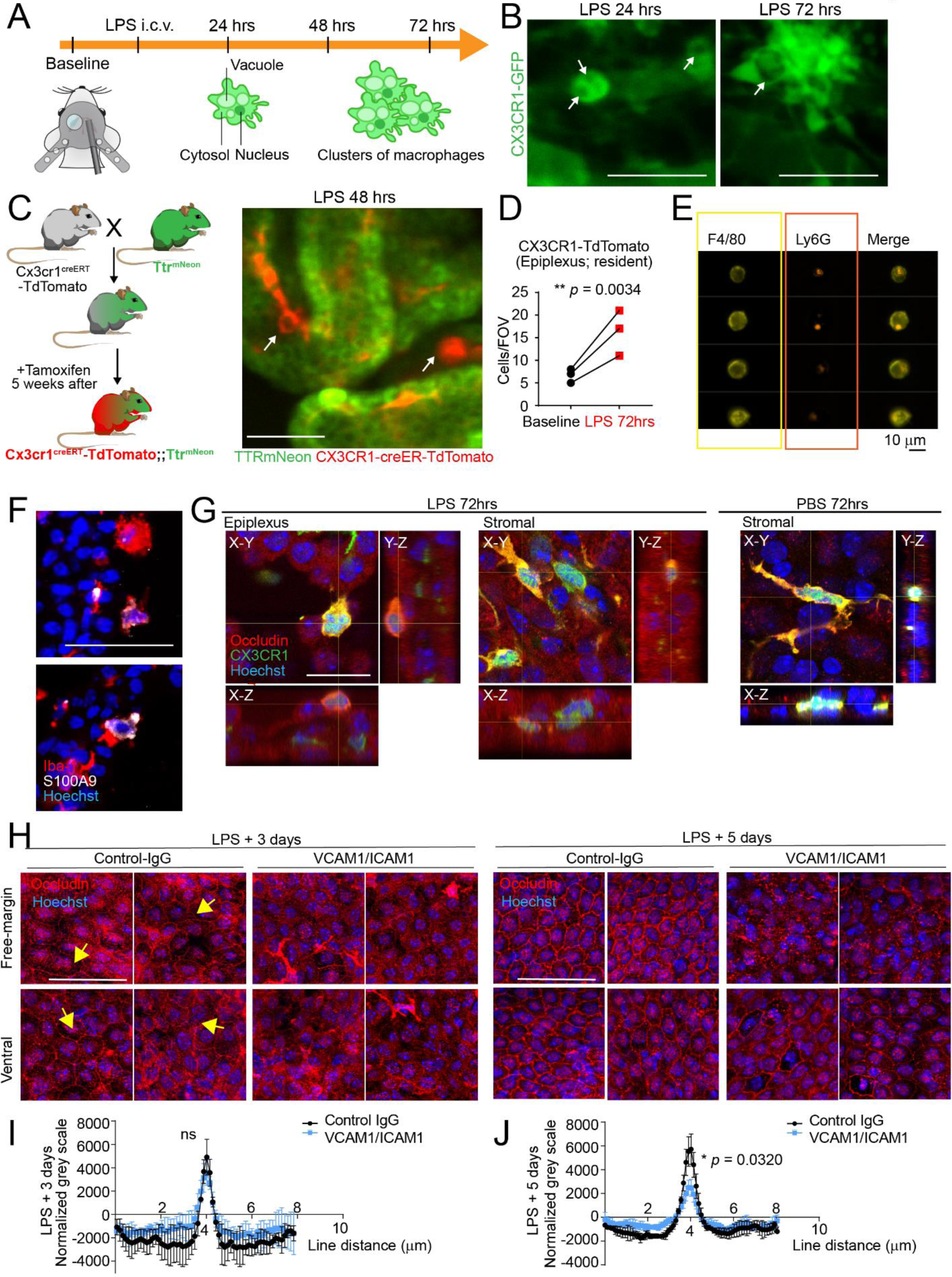
Macrophages at the ChP aided inflammation resolution and barrier repair. (**A**) Schematics depicting morphological changes of ChP macrophages following LPS ICV. (**B**) Representative images extracted from *in vivo* imaging showing ChP macrophages with large vacuoles and forming clusters. Also see Supplemental video 9-10. Scale = 50 µm. (**C**) Schematics demonstrating the breeding scheme to acquire mice harboring TTR^mNeon^ that illuminates epithelial cells and expressing Tamoxifen-inducible Cx3cr1^Tdtomato^ that labels resident macrophages. The image on the right is a representative image extracted from *in vivo* imaging, demonstrating morphological changes in epiplexus resident ChP macrophages. Also see Supplemental video 11. Scale = 50 µm. (**D**) Quantification showing significant increase of CX3CR1+ resident epiplexus macrophages from baseline to 72 hours following LPS ICV delivery (** *p* = 0.0034, N=3). (**E**) Representative images from ImageStream showing Ly6G+ neutrophils internalized by F4/80+ macrophages. Scale = 10 µm. (**F**) Representative images showing co-localization of Iba-1 (macrophage) and S100A9 (neutrophil) by immunohistochemistry. Scale = 50 µm; (**G**) Representative 3D image and orthogonal views showing epiplexus and stromal CX3CR1+ macrophages that contain Occludin in ChP explants from mice 72 hours following LPS vs. PBS ICV delivery. (**H-J**) Representative images and line profile quantification showing progressive repair of ChP epithelial tight junctions. Mice treated with anti-VCAM1/ICAM1 neutralizing antibodies (N=4) had delayed Occludin repair than controls (N=3). Statistical significance was calculated by the maximum height of peaks. * p = 0.0320. Scale = 50 µm. All quantitative data are presented as mean ± SD.

Next, we questioned the roles of gained epiplexus macrophages in restoring epithelial cell junctions following LPS-induced breakdown. Occludin was detected within CX3CR1+ immune cells in both LPS-treated mice and PBS controls, but LPS-treated mice had many Occludin+ epiplexus immune cells while PBS control mice had few Occludin+ macrophages and these were predominantly located within the stromal space (**Figure 6G**, more examples in **Figure S9D**). This finding suggests that, while stromal macrophages may be involved in barrier maintenance at baseline, epiplexus macrophages might be involved in epithelial barrier repair following LPS ICV, possibly by cleaning up epithelial debris which is a commonly known function of macrophages during wound healing. We found that male mice treated with VCAM1/ICAM1 neutralizing antibodies 24 hours following LPS to reduce epiplexus macrophages had delayed recovery of Occludin over 5 days compared to their IgG-treated controls, as reflected by weakened signal peaks between two adjacent cells indicating incomplete barriers (**Figure 6H-J**). In female mice, the recovery rate varied across all conditions, and the differences were less noticeable (tested with N=8 in two independent experiments). We conclude that the recruited ChP epiplexus macrophages by apical VCAM1/ICAM1 aided in the repair of epithelial junctions which contributes to re-establishing the blood-CSF barrier after inflammation.

## DISCUSSION

The ChP has long captivated interest as a gateway for peripheral immune cell entry into the brain [1]. However, definitive evidence demonstrating immune cell extravasation across the ChP brain barrier has been lacking due to challenges inherent to studying the ChP barrier in real-time, deep inside the brain. Here, we apply recent advances in chronic *in vivo* two-photon imaging [41] along with transcriptomics, lineage tracing, and histology to demonstrate that the ChP coordinates a systematic, multi-cellular response in response to CNS inflammation in a mouse model of meningitis (**Figure 7**). Longitudinal live imaging revealed that the ChP organized immune recruitment involving both peripherally- and centrally (CSF)-sourced myeloid cells. When we combined these observations with single-cell transcriptomics and histological approaches, a molecular map emerged revealing ChP epithelial cells as key orchestrators overseeing the ChP immune response. We uncovered cell-cell interactions that spanned (1) immune cell recruitment (upregulation of cytokines and chemokines), (2) barrier breakdown (upregulation of matrix metalloproteases), (3) immune cell survival (upregulation of colony-stimulating-factor CSF1), and (4) immune cell adhesion to the ChP (upregulation of VCAM-1 and ICAM-1). Collectively, this series of steps culminated in the clearance of neutrophils/monocytes, ChP barrier repair, and resolution of inflammation by recruited macrophages. These findings highlight the ChP as an immune organ that deftly coordinates both local and long-range signaling to ward off inflammation.

**Figure 7.**
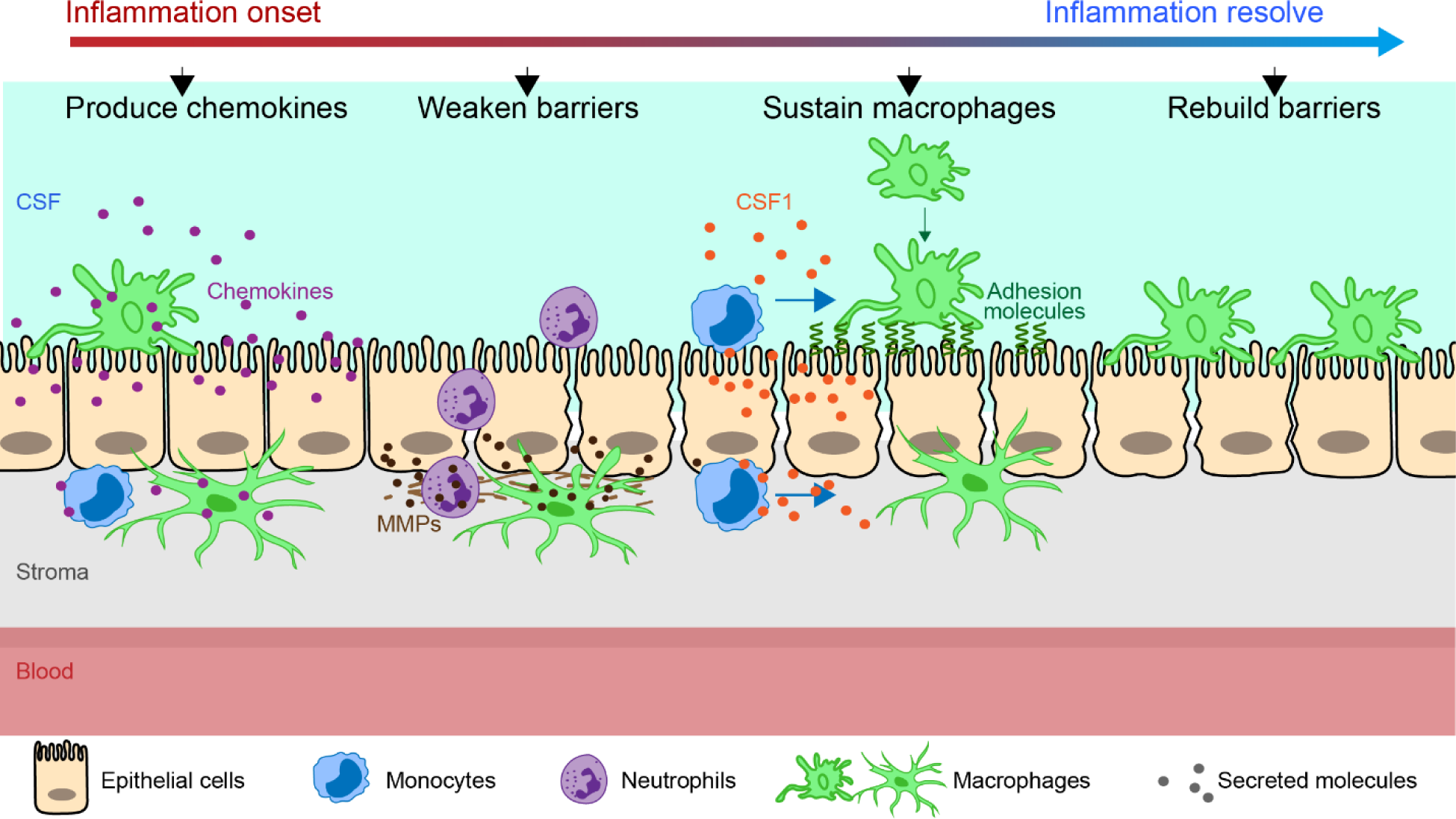
Summary schematics depicting the multitude of epithelial-immune interactions during ChP inflammation. From the onset to the resolution of inflammation, the ChP epithelial cells and immune cells collaborated to (from left to right): (1) produce chemokines to recruit immune cells to the ChP; (2) weaken the epithelial barrier through extracellular matrix remodeling to facilitate infiltration; (3) produce adhesion molecules (ICAM1 and VCAM1) and key differentiation and survival factors (colony-stimulating factor CSF1) to sustain macrophages, which (4) contributed to rebuild the epithelial barrier.

Our data demonstrate that ChP epithelial cells provide a critical immune cell niche, thereby extending the concept of the stem cell niche that the ChP-CSF system provides for the developing and adult [56, 57], we discovered a transiently inflamed ChP epithelial cell population (Inf-Epi, **Figure 5**) with upregulated expression of colony-stimulating-factor CSF1, a critical survival and differentiation signal for macrophages [54], as well as increased section of inflammatory chemokines. The same epithelial population also produced the extracellular matrix remodeling proteins to promote leukocyte infiltration and the adhesion molecules that facilitate CSF-to-ChP macrophage recruitment for barrier repair following inflammation. Thus, these inflamed epithelial cells ultimately provide key signals needed to coordinate the recruitment, adhesion, barrier breakdown, and eventual repair of the barrier. These data establish the concept that ChP epithelial cells essentially function as adaptable “general contractors,” remodeling the barrier as needed to address the task at hand.

The source (peripheral or central), type of pathogen (bacterial, viral, parasite), and extent of inflammation (acute or chronic) likely determine the molecular mechanisms involved and the extent of the ChP inflammatory response. Early studies of ChP inflammation focused primarily on T cells [10, 25, 30, 58]. In our studies, we did not detect a robust T cell response. Rather, informed by the cellular composition of CSF in our meningitis model mice, we focused our analyses on neutrophils and macrophages for their roles in anti-bacterial activities and tissue remodeling. ChP macrophage responses in our meningitis model differ from models of peripheral inflammation, including other conditions involving LPS [41, 59–61]. For example, our prior work showed that peripheral LPS delivery triggered an elongation of stromal ChP macrophages along blood vessels [41], which likely reflects an adaptable extra barrier as a means of protection against peripheral insults. A similar process was recently reported by microglia following blood vessel injury in the brain [39].

The function(s) of ventricle-facing ChP macrophages has remained elusive since their initial description over 100 years ago [10]. Live imaging recently captured these cells as immediate responders to micro-hemorrhage at the ChP [41]. Here, we discovered that the ChP can expand this cell population by both recruiting monocytes from the blood and attracting macrophages from the CSF, akin to organ-cavity recruitment in peripheral organs [38]. The source of these CSF macrophages remains uncertain, as scRNAseq suggests these cells lack signature genes (e.g., *Sall1*) of either microglia or border-associated macrophages (BAMs). As transcription in microglia and macrophages is highly dynamic, it is possible that CSF macrophages represent either mobilized microglia and BAMs, or newly differentiated macrophages from infiltrated monocytes that travel in between the ChP and the ventricle wall, or a combination of cells recruited from multiple sources. Despite their exact origin, we found that these recruited epiplexus macrophages attached to the ChP surface through surface adhesion molecules such as ICAM1 and VCAM1. We, and others [12, 55], have shown that these adhesion molecules are upregulated on the CSF-facing surface of ChP epithelial cells, but not in other tissues and brain vasculature. Here, we provide the first *in vivo* evidence demonstrating the significance of this unconventional expression pattern in retaining epiplexus macrophages at the ChP, including those recruited to the ChP from the CSF. Preventing macrophage attachment to the ChP by neutralizing VCAM1 and ICAM1 hindered the recovery of ChP epithelial tight junctions. Notably, the phenotype was more distinctive in males compared to females. Together, these data suggest that these very same ChP macrophages that respond rapidly to acute tissue injury also participate in brain barrier repair, thereby assigning a critical second function for these cells.

In the forebrain, the ChP extends up into the lateral ventricles from its brain attachment site ventral to the hippocampus and fornix. Due to limitations in the commonly available imaging optics, our imaging preparation captures approximately the top one third of the ChP. A noteworthy challenge moving forward will be to adapt newly available technologies (e.g., three-photon imaging [62]) to reach deeper into the ventricles to track the path of immune cells in real time near the base of the ChP. For example, our histological analyses (**Figures S1J and 2I**) also revealed leukocytes accumulating in the stroma at the “base” of the ChP where it connects to the brain parenchyma, but we were not able to obtain live imaging data of leukocyte movements and entry into the ChP in this region. A broader span of imaging depths will eventually also allow tracking of cells as they travel beyond the currently imaged field of view. Ultimately, a toolbox consisting of mouse models, fluorescent reporter lines, live imaging, lineage tracing, cell labeling, and omics approaches will help unravel the dynamics of all immune cell types at the ChP (including T cells and B cells that are not the focus of this particular study).

Leukocyte recruitment to activated tissues is well-established, and prevailing models posit that peripheral monocytes are recruited from blood to produce new macrophages for organ barrier repair [63]. Recent work in traumatic brain injury also demonstrates leukocyte infiltration through broken blood vessels and identifies specialized microglia that rebuild the vessels and repair the blood-brain barrier [39]. Our work focused on the ChP as both an immune organ that responds to immune challenges and a unique immune barrier that actively attends to only to its own needs but to the brain’s immune needs. However, given the complex immune environment of the brain and its many barriers (e.g., the meningeal barrier, the blood-brain barrier, and circumventricular organs [1]), the ChP likely collaborates with other brain compartments to ward off inflammation.

In summary, our work reveals that the ChP is an active immune organ which responds to inflammation with complex immune trafficking and timely barrier repair through elaborate epithelial-immune crosstalk and collaboration. Considering the growing number of neurologic diseases associated with ChP immunity, our studies open up new avenues for considering how the ChP brain barrier can be activated and harnessed to better understand and potentially treat neuroinflammation.

## Supporting information

MATLAB code

## ACKNOWLEDGEMENTS

We thank members of the Lehtinen, Ordovas-Montanes, Chiu, and Andermann labs, Denisa Wagner, Tanya Mayadas-Norton, and Michael Carroll for helpful discussions, and Nancy Chamberlin for critical reading and editing of the manuscript. We thank the following facilities and personnel: Chinfei Chen, Hisashi Umemori, Cheng-Hao Chien and the BCH IDDRC Cellular Imaging Core; Michael Anderson, Bin Bao, and the Cell Function and Imaging Core (Harvard Digestive Diseases Center); BCH: PCMM flow cytometry facility; BCH pathology core; and the Harvard Medical School Electron Microscopy Facility. This work was supported by: BrightFocus postdoctoral fellowship in Alzheimer’s Disease research (H.X.); HHMI James H. Gilliam Fellowships for Advanced Study program (P.L.). Edward R. and Anne G. Lefler Center Postdoctoral Fellowship and Hebrew University Postdoctoral Fellowship (S.G.); T32 NS007473-22 (C.H.); T32 GM007753, T32 GM144273, and American Heart Association Pre-doctoral Fellowship (M.E.Z.); NSF Graduate Research Fellowship (F.B.S.); Harvard College Research Program (M.T.); Burroughs Wellcome Fund, Chan-Zuckerberg Initiative, and NIH R01 AI168005 (I.M.C.); NIH Pioneer Award DP1 AT010971 (M.L.A.); The Pew Charitable Trusts Biomedical Scholars, NIH R01 HL162642, and The Cell Discovery Network at Boston Children’s Hospital supported by the Manton Foundation and Warren Alpert Foundation (J.O.M.); The New York Stem Cell Foundation (M.K.L. and J.O.M.); Cure Alzheimer’s Fund, Human Frontier Science Program (HFSP) research program grant #RGP0063/2018, and NIH R01 NS088566, RF1048790 (M.K.L.); BCH IDDRC 1U54HD090255. The content is solely the responsibility of the authors and does not necessarily represent the official views of the NIH.

## AUTHOR CONTRIBUTIONS

H.X. and M.K.L. conceptualized and designed the study; H.X., P.L., S.G., F.S., J.C., J.O., M.L.A. and M.K.L. established methodology; H.X., S.G., A.P., M.E.Z., J.C., M.W., and L.D. conducted experiments; H.X., P.L., S.G., F.B.S., C.H., J.C., M.T., M.E.Z., M.L.A., J.O., and M.K.L. analyzed data; L.D., H.G.W.L., and I.C. provided experimental material; I.C., J.O., M.K.L. provided funding; M.K.L. supervised the study; H.X., P.L., J.O., and M.K.L. wrote the manuscript. All co-authors read and approved the manuscript.

## DATA AVAILABILITY

Sequencing data (Figure 1, Figure 4, Figure S1, and Figure S3) will be available in GEO upon publication. All other data are available from the authors. All biological material were either directly commercially available or are available upon request from the lab.

## CONFLICT OF INTERESTS

J.O.M. reports compensation for consulting services with Cellarity, Tessel Biosciences, and Radera Biotherapeutics.

## METHODS

### Animals

The Boston Children’s Hospital IACUC approved all experiments involving mice in this study. The following genetic strains were obtained from JAX laboratory: Cx3cr1GFP (JAX 005582), Ccr2RFP (017586), Cx3cr1CreER (020940), Ai6 (007906, ZsGreen reporter), Ai14 (007914, TdTomato reporter), Ly2Cre (004781). C57BL6 mice were obtained from Charles River Laboratories. C57BL6-TTRmNeonGreen was created with Boston Children’s gene manipulation core (referred to as TTRmNeon in the manuscript) [48]. Both male and female mice were equally included in the study except for single cell sequencing and CSF cell transplanting assays, where only male mice were used. Animals were housed in a temperature-controlled room on a 12-hr light/12-hr dark cycle and had free access to food and water.

### Human samples

All human samples were obtained using an IRB-approved protocol at Boston Children’s Hospital. Three specimens with diagnosed bacterial meningitis (patients: 18-year-old female, 10-day-old male, and 2-week-old male) and two non-infectious control cases were analyzed by immunohistochemistry. Images were acquired using an Olympus brightfield microscope with DP71 camera and cellSens Entry software.

### Intracerebroventricular injection (ICV)

Adult mice were anesthetized using isoflurane vaporizer. An incision was made along the midline of scalp to expose the skull. The lateral ventricle was located by stereotaxic coordinates (bregma −0.4mm, midline +1mm, dura −2.2mm). A single delivery path was prepared above the lateral ventricle. 5 µl of lipopolysaccharide solution (0.5mg/ml) was delivered by a Hamilton syringe to the lateral ventricle over the course of 5 min.

The following reagents were delivered by ICV: Lipopolysaccharides (Sigma L3024, 0.5mg/ml x 5 µl). Rat Ultra-LEAF™ purified anti-mouse CD106 antibody, clone 429 (MVCAM.A) (BioLegend 105728, 1µg/mouse), rat Ultra-LEAF™ purified anti-mouse CD54 antibody, clone YN1/1.7.4 (for ICAM1, BioLegend 116133, 1µg/mouse). Rat purified anti-mouse CD11a antibody (For LFA-1α, BioLegend 101101, 1µg/mouse), rat purified anti-mouse CD49d antibody (For VLA-4α, BioLegend 103701, 1µg/mouse), rat IgG2b κ control antibody (ThermoFisher Scientific, 17-4031-82, matching dose to other antibodies).

In addition, live and heat-killed *S. agalactiae* (GBS) were delivered by ICV. All procedures related to culture and work with pathogenic bacteria were approved by the Committee on Microbiological Safety of Harvard Medical School and conducted under Biosafety Level 2 guidelines. GBS clinical isolate COH1 (serotype III) was grown in Todd-Hewitt broth (THB)(Sigma) at 37° C and growth was monitored by measuring optical density at 600 nm (OD600). To prepare bacteria for intracerebral ventricular injection, GBS was grown to mid-log phase, pelleted by centrifugation, and resuspended in PBS (Sigma). Heat-killed bacteria were prepared by boiling the bacterial suspension for 5 min. THB was inoculated with a portion of the heat-killed suspension and incubated at 37° C for 2 days to confirm lack of bacterial growth.

### CSF collection and analysis

CSF was collected by inserting a glass capillary into cisterna magna, and collected CSF was centrifuged at 1000xg for 10min. at 4° C to remove any tissue debris. The supernatants were used to measure cytokines and chemokines using LEGENDplex™ Mouse Inflammation Panel (13-plex) (BioLegend 740446). In the case of CSF cell transplant, the samples were centrifuged in the same way, and the pellet was resuspended in sterile PBS for subsequent analysis.

### Tissue collection and processing: explants and brain blocks

Animals were anesthetized by ketamine and perfused first with ice cold PBS, and then cold 4% paraformaldehyde (PFA). For cryosectioning, brains were dissected and further fixed in 4% PFA at 4° C overnight, and then incubated in 30% sucrose for at least 48 hours, followed by OCT (1 hour on ice) prior to freezing by dry ice and 2-met-butane bath. For wholemount ChP explant, the LVChPs were dissected and incubated in 4% PFA at room temperature for 5 min., and then proceed to immunostaining.

### Immunostaining

Cryosections and explants were blocked and permeabilized (0.3% Triton-X-100 in PBS; 5% serum), incubated in primary antibodies overnight and secondary antibodies for 2 hours. Sections and explants were counterstained with Hoechst 33342 (Invitrogen H3570, 1:10,000) and mounted using Fluoromount-G (SouthernBiotech).

The following primary antibodies were used: chicken anti-GFP (Abcam ab13970; 1:1000), rabbit anti-RFP (Rockland 600-401-379, 1:500), rat anti-PECAM (BD Pharmingen 550274, 1:100), rat anti-CD45 (Fisher Scientific BDB550539, 1:50), goat anti-S100A9 (R&D Systems AF2065, 1:200), rabbit anti-Occludin (ThermoFisher 71-1500, 1:50), rabbit anti-Iba1 (Wako 019-19741, 1:200), goat Anti-Type IV Collagen-Alexa Fluor® 488 (Southern Biotech 1340-30, 1:200), PE rat anti-mouse CD62E (for E-selectin, BD Biosciences 553751, 10µg per mouse), Alexa594 anti-mouse Ly-6G/Ly-6C (Gr-1) antibody (BioLegend 108448, 10ug per mouse), rat Ultra-LEAF™ purified anti-mouse CD106 antibody, clone 429 (MVCAM.A) (BioLegend 105728, 1:50), rat Ultra-LEAF™ purified anti-mouse CD54 antibody, clone YN1/1.7.4 (for ICAM1, BioLegend 116133, 1:50), rat purified anti-mouse CD11a antibody (For LFA-1α, BioLegend 101101, 1:50), rat purified anti-mouse CD49d antibody (For VLA-4α, BioLegend 103701, 1:50), rabbit anti-claudin2 (Thermo-Fisher, 51-6100, 1:100), rabbit anti-ZO1 (Thermo-Fisher, 61-7300, 1:100). Goat anti-mouse MCSF antibody (for CSF1, R&D AF416, 1:100). Secondary antibodies were selected from the Alexa series (Invitrogen, 1:500). Images were acquired using Zeiss LSM880 confocal microscope.

### Confocal microscopy and image analysis

All confocal images were acquired using Zeiss LSM 880 with Airyscan FAST confocal microscope. Explant images were taken with 2.5µm z-stacks. All images were processed with FIJI [64]. CX3CR1-CCR2 colocalization quantification (**Figure 3D**) was performed with MATLAB (2020b). The code is included as a Supplemental file (ChP_red_green_Colocalization.m)

### Intracardiac injection (IC)

Adult mice were anesthetized using isoflurane vaporizer. The hair on the left chest was removed. An insulin syringe was used to deliver 200 µl total of solution directly into the left ventricle of heart. Alexa594 anti-mouse Ly-6G/Ly-6C (Gr-1) antibody (BioLegend 108448, 10ug per mouse) and PE rat anti-mouse CD62E (for E-selectin, BD Biosciences 553751, 10µg per mouse) were delivered with this approach for live *in vivo* labeling. Anti-VCAM1 and anti-ICAM1 antibodies were delivered IC to block endothelial expression. LPS (2.5 µg) was delivered IC to compare to ICV.

### Headpost, cranial window, and ICV canula placement

Mice used for *in vivo* two-photon imaging (4-6 months) were outfitted with a headpost, a 3 mm cranial window over the left lateral ventricle, and a contralateral trans-occipital cannula for ICV injections as previously described [65]. A guide cannula was inserted and held in place with a screwcap to prevent infection and tissue growth into the cannula. This guide cannula was removed prior to ICV injections.

### Two-photon imaging and image processing

Two-photon microscopy (Olympus FVMPE-RS two-photon microscope; 512 x 512 pixels / frame) was used to record immune cells activities in ChP in vivo from the following mice: Cx3cr1GFP (JAX 005582), Ly2Cre (004781) crossed with Ai6, *TTR^mNeon^* crossed with Cx3cr1CreER (020940) and Ai14 reporter line. 200µm-400µm Z-stack was acquired with 5µm stepping size at each field of view over the course of approximately 1 hour. Imaging and analyses were performed as described [41]. A 25X magnification, 8 mm working distance objective was used.

LPS solution (0.5mg/ml) was delivered to the lateral ventricle of live mice prior imaging through the injection cannula at a rate of 1 μL / min over 5 min.

Images were registered and processed following the established algorithm [41]. MATLAB code used to match two-photon in vivo views with post hoc explant images is included in the Supplemental files (PostHocMatching.m).

### Tamoxifen induction of gene expression

Tamoxifen was dissolved in canola oil at 20mg/ml concentration. The solution was incubated at 37° C overnight with shaking, and stored at 4° C with light protection. Mice were injected intraperitoneally (i.p.) daily for 4 days, 100µl per day. All mice with receiving tamoxifen treatment were subjected to experiments at least 5 weeks from the last dose.

### Blood cell transfusion

Lyz2-ZsGreen mice and WT mice both received LPS ICV. 18 hours later, blood was collected from Lyz2-ZsGreen mice, treated with citric acid to prevent clotting, and incubated with 10x volume of ACK lysis buffer to remove red blood cells. The remaining cells were pelleted and washed, and resuspended in sterile PBS. The cell suspension was injected to WT mice by IC. WT mice were harvested 6 hours after cell transfusion for brain histology.

### ImageStream

Freshly dissected LV and 4V ChP were placed in digestion solution (HBSS + collagenase/dispase, Sigma-Aldrich #10269638001) for 30 mins at 37° C with shaking at 600rpm. Then, DNase1 was added (4 µl for 100 µl digestion solution) and incubated for 15 mins. 1ml DMEM was added to stop enzyme activity, and cells were collected by centrifuging (10 mins at 400g, 4° C). The cells were first stained for outer markers with KIRAVIA 1:100 (BioLegend 405172) + F4/80-PE 1:100 (BioLegend, #157340) in 0.5%BSA/HBSS for 25 min. on ice, and then fixed with Cytofix/Cytoperm (BD bioscience, cat # 554714, 50 µl for sample for 20 min. at room temperature). Next, cells were stained for inner marker with Ly6G-AF594 (BioLegend 108448) 1:100 for 20 min. at room temperature and incubated with Hoechst 1:2000 (Sigma # 23491-45-4) prior to analysis with ImageStream (Flow Cytometry PCMM core facility at BCH and Harvard Medical School).

### Electron microscopy

All tissue processing, sectioning, and imaging was carried out at the Conventional Electron Microscopy Facility at Harvard Medical School. The ChP were fixed in 2.5% Glutaraldehyde/2% Paraformaldehyde in 0.4% CaCl2 and 0.1 M sodium cacodylate buffer (pH 7.4). They were then washed in 0.1M cacodylate buffer and postfixed with 1% Osmiumtetroxide (OsO4)/1.5% Potassiumferrocyanide (KFeCN6) for one hour, washed in water three times and incubated in 1% aqueous uranyl acetate for one hour. This was followed by two washes in water and subsequent dehydration in grades of alcohol (10 minutes each; 50%, 70%, 90%, 2×10min 100%). Samples were then incubated in propyleneoxide for one hour and infiltrated overnight in a 1:1 mixture of propyleneoxide and TAAB Epon (Marivac Canada Inc. St. Laurent, Canada). The following day, the samples were embedded in TAAB Epon and polymerized at 60° C for 48 hours. Ultrathin sections (about 80nm) were cut on a Reichert Ultracut-S microtome, and picked up onto copper grids stained with lead citrate. Sections were examined in a JEOL 1200EX Transmission electron microscope or a TecnaiG² Spirit BioTWIN. Images were recorded with an AMT 2k CCD camera.

### Tissue dissociation for single cell sequencing

The lateral ventricle (LV) ChP was dissected in cold RPMI media with 10mM HEPES and minced with a pair of fine scissors inside a 1.5 mL tube (roughly 200 times until only small pieces remain). LV ChP from 3 adult male mice were pooled into each sample. Tissue pellets were digested in an enzyme cocktail containing collagenase P (0.5 mg/mL, Sigma cat. 11213865001), dispase (0.8 mg/mL, Worthington cat. LS02104), and DNAse1 (250 U/mL, Worthington cat. LK003172) for 30min. at 37° C with slow head-to-toe rotation. Post-digestion tissues were washed, triturated, and filtered with 70µm mesh. All pipette tips and tubes were coated by incubating in HBSS containing 2% BSA. To avoid transcriptional changes during the dissociation process, dissection and digestion solutions contained actinomycin (Sigma, cat. A1410, 5 µg/ml), triptolide (Sigma, cat. T3652, 10µM), and Anisomycin (Sigma, cat. A9789, 27.1 µg/ml) [66]. For CSF analysis, CSF from 5 adult male mice were collected and pooled into each sample. The CSF was treated with ACK lysing buffer (Thermo Fisher A1049201) to remove red blood cells. Cells from CSF were pelleted at 1000g x 10min (4° C) and washed.

### Single cell sequencing

Single cell suspension from digested ChP or CSF was evaluated under microscope with trypan blue staining for cell viability and cell number. In ChP study and CSF study respectively, 0.5-1 million live cells from each condition were hashed by TotalSeqTM B0301, B0302, and B0303 anti-mouse Hashtag antibodies which target MHCII and CD45 (BioLegend, cat. 155831, 155833, 155835), and pooled at equal ratios. Cells were counted again after hashing and adjusted to 1500 live cells per µl. Approximately 20,000 cells per sample were loaded onto the Chromium controller. Libraries were prepared according to manufacturer protocols (Chromium Next GEM Single Cell 3’ Reagent Kits v3.1 (Dual Index)). Sequencing was performed at Broad Institute (NovaSeq SP) with ∼20,000 reads/cell depth.

### Single cell data Preprocessing and quality control

Gene expression and feature barcoding data were processed using CellRanger (v3.1.0) with alignment to the mm10 reference genome. To assign ChP cells to different conditions, feature barcodes (TotalSeq) were demultiplexed using the HTODemux() function in Seurat v4 with a quantile threshold of 0.99, yielding 7,987 singlets, 1,876 doublets, and 10,412 unassigned cells. Doublets and low-quality cells with greater than 30% of reads mapping to mitochondrial genes were removed. To assign CSF cells to either 24 hr or 48 hr, feature barcodes were demultiplexed similarly, yielding 11,770 singlets, 1,270 doublet, and 27 unassigned cells. Doublets and cells with greater than 10% of reads mapping to mitochondrial genes were removed.

#### Cell Clustering and Annotation

Dimensional reduction, cell clustering, and differential gene expression analysis were conducted using Seurat v4.2.1 [67]. We filtered cells that were identified as likely doublets through hash demultiplexing then gene expression was scaled and normalized using SCTransform [68, 69] with method set to “glmGamPoi” [70] and dimensionality reduction was conducted using principal component analysis (PCA). For CSF cells, we used 31 PCs for dimensional reduction and clustering at resolution of 0.5. For ChP cells, we selected PCs visually using an elbow plot then used these PCs for cell clustering at high resolution to further identify likely doublets and contaminating erythrocytes based on high levels of co-expression of marker genes from disparate classes of cell types. We iteratively removed these cells and re-clustered until our analysis yielded a dataset of 16,294 cells. ChP cells wereassigned cell type identities based on expression of known canonical marker genes, including epithelial (Kcnj13, Wfdc2, and Ecrg4), fibroblast (Dcn, Alpl, and Igfbp4), myofibroblast (Acta2, Myl9, and Tagln), pericyte (Pdgfrb, Cox4i2, and Kcnj8), endothelial (Ly6a, Kdr, and Flt1), myeloid (Ms4a6c, Cx3cr1, and Trem2), neutrophil (Csf3r, Il1b, and S100a8), and lymphocyte (Ptprcap, Trbc2, and Nkg7). For detailed annotation of cell subtypes and states, we subclustered each cell type individually and annotated cells based on their gene expression profiles and known markers.

#### Cell-Cell Communication Analysis using CellChat

We used CellChat (v1.5) [71] to infer cell-cell communication based on the expression of ligand-receptor pairs and downstream signaling mediators from our scRNAseq data. We loaded in normalized counts and applied standard parameters to assess signaling between cell types. For comparison of signaling between different timepoints, we collapsed cell states into higher-level clusters (e.g. Resident Macrophages and Inflammatory Resident Macrophages were combined into “Resident Macrophages”) and ran CellChat with multiple comparisons.

#### Packages used

Seurat (v4.2.1 [67])

CellChat (v1.5 [71])

Circlize (v0.4.16 [72])

ComplexHeatmap (v2.13.4 [73])

Ggplot2 (v3.4)

scCustomize (v0.7.0 [75])

### PLX5622 treatment

PLX5622 chow were manufactured by ScottPharma at 1200ppm, using regular chow provided at Boston Children’s Hospital as contorl. PLX-5622 compound was provided by Cayman Chemical (#28927). Mice were placed on PLX5622 diet or control diet for 10 days before LPS ICV, and kept on the same diet until tissue harvest.

### CSF cell transplant

CX3CR1-ZsGreen mice were injected ICV with LPS as described. 48 hours following LPS injection, CSF was collected as described. Cells within CSF were collected by centrifuging the CSF at 1000xg for 10 min at 4° C, and washed once with cold sterile PBS. The cells were resuspended in PBS for transplant at 10 µl per donor mouse (i.e. When 5 mice were used as donors, the CSF cells were resuspended to 50µl). 5 µl of cell suspension was injected ICV to WT recipient mice that received LPS ICV 24 hours before. The recipient mice were treated by ICV injection of 1µg each VCAM1/ICAM1 antibody cocktail or control rat IgG 10 min. before ICV injection of cell suspension. Recipient mice were harvested 48 hours after cell transplant.

### Quantification of Occludin

Line profile analysis from FIJI was used to quantify the strength and distribution of Occludin between adjacent cells (approach was modified from [74]). 3 images from each explant were used for analysis. A total of 20 adjacent cell pairs were selected based on nuclei positions. A line of 8 µm was drawn between the two nuclei and registered as ROI. The same line was then placed on the Occludin image. Adjustments in line positions were made so the middle point of the line was located to the Occludin barrier between the cells, unless Occludin signal was completely missing, in which case the line was placed evenly in between the 2 nuclei. Line profile was analyzed to determine the peak signal at the cell border (4 µm distance). The average height of all 20 peaks were used to represent each mouse. Peak height was used for statistical analysis between conditions.

## Supplemental figures

**Supplemental figure 1.**
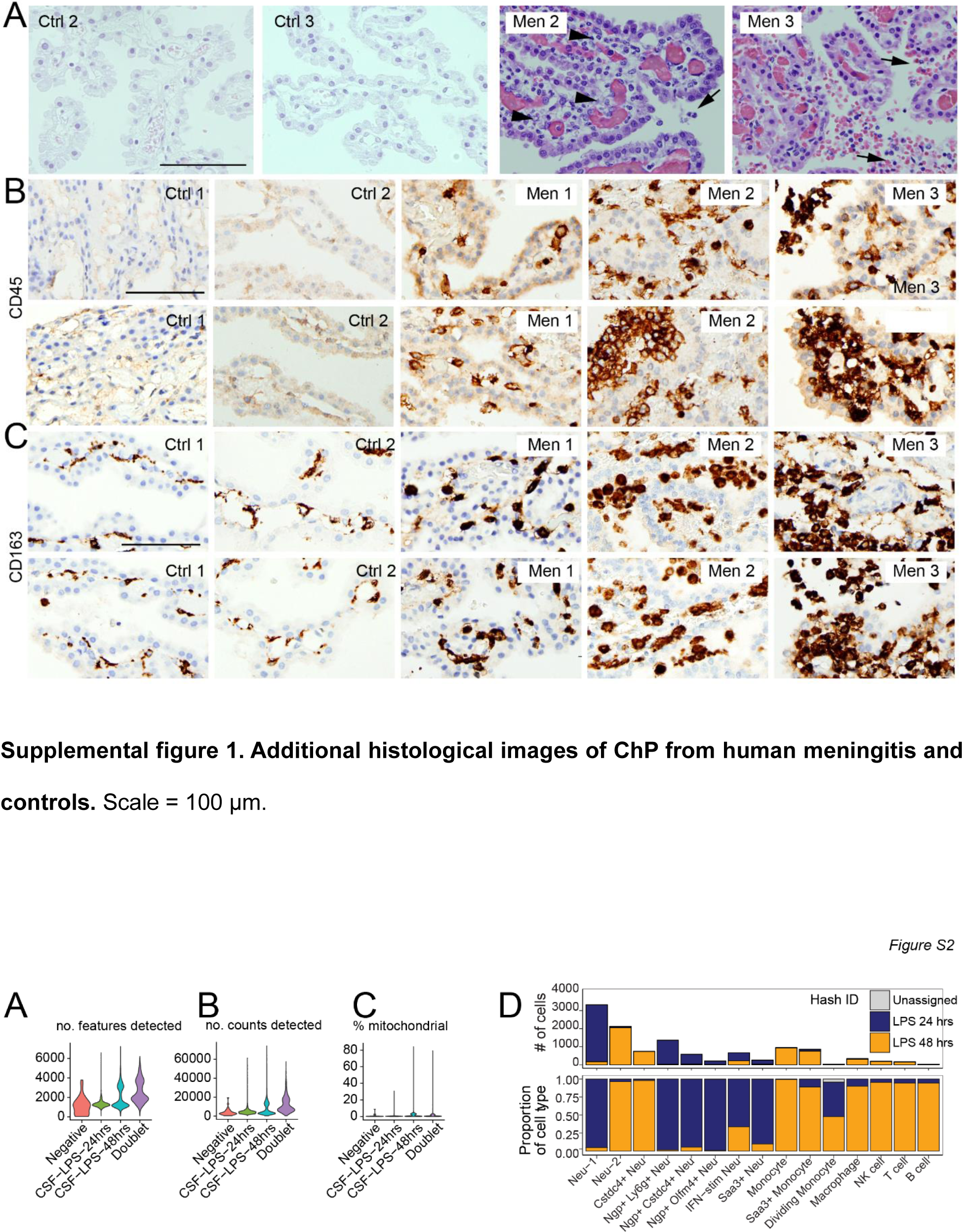
Additional histological images of ChP from human meningitis and controls. Scale = 100 µm.

**Supplemental figure 2.**
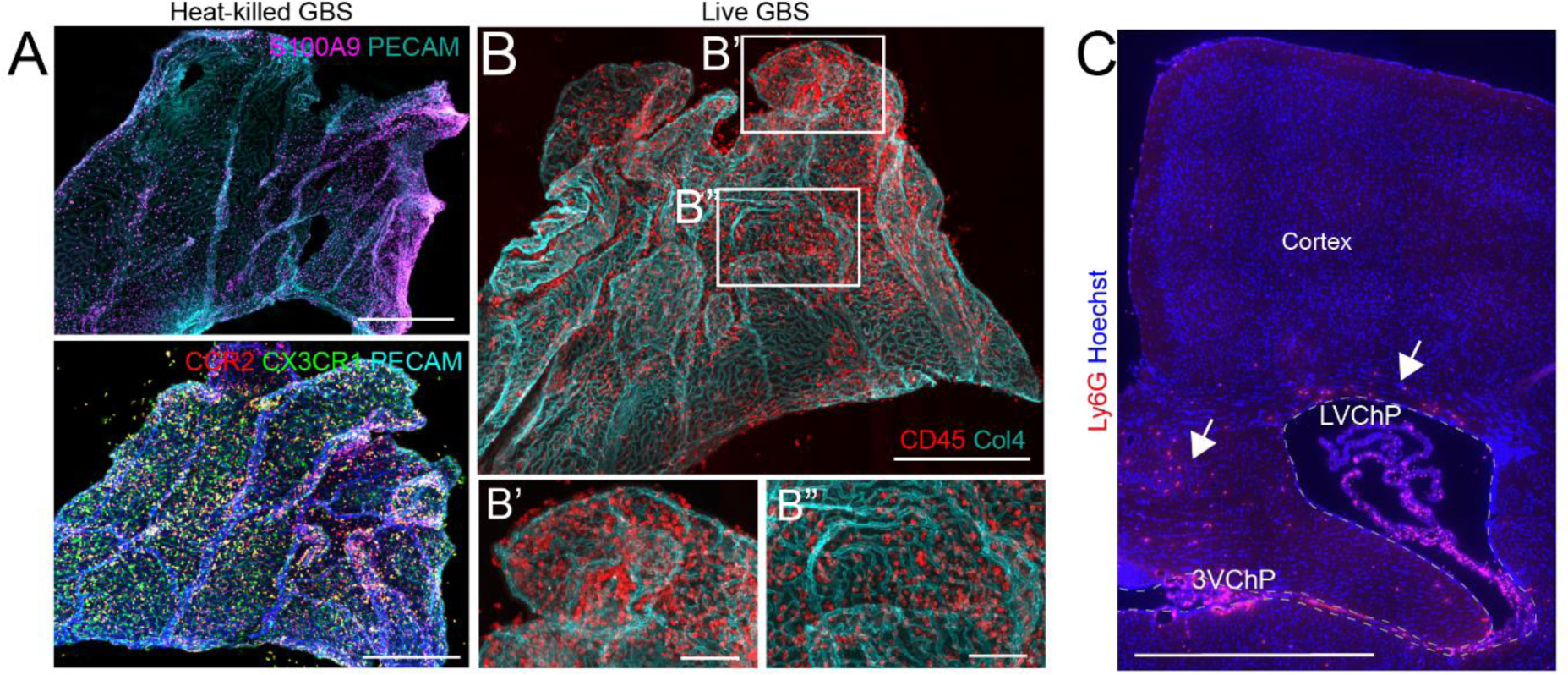
Supportive data of CSF scRNAseq. (**A-C**) QC plots from CSF scRNA-seq showing comparable gene detection, cell count, and cell viability across samples. (**D**) Cell frequencies and relative proportions by assigned has identity.

**Supplemental figure 3.**
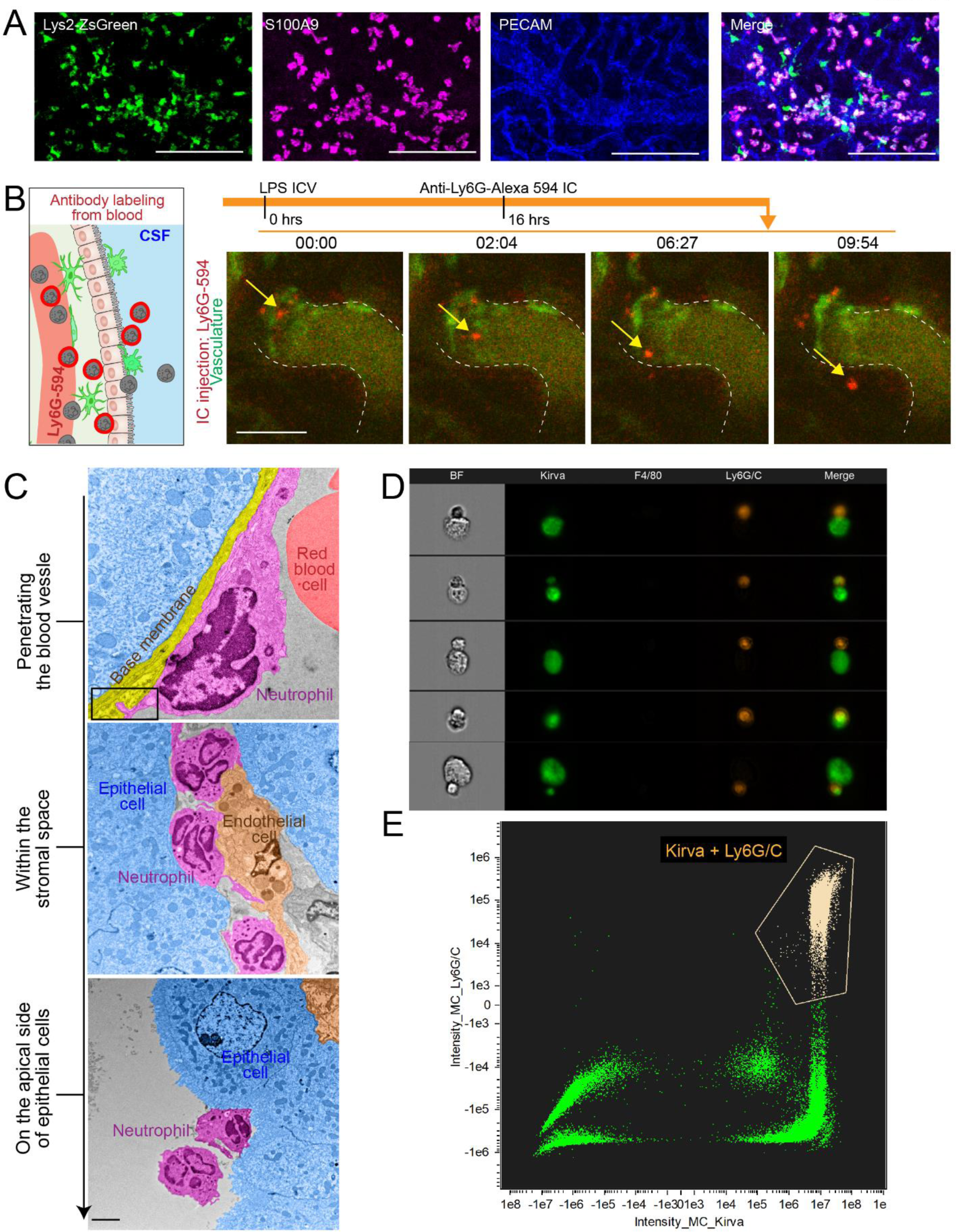
Additional images showing neutrophils and monocytes infiltrate through the ChP following brain infection. (**A-B**) Representative images of wholemount ChP explant showing neutrophils following heat-killed Group B Strep (GBS) ICV (H) and live GBS ICV (I). Scale = 500 µm. (**C**) Representative image of brain coronal section showing neutrophils, marked by Ly6G, predominantly accumulated in the 3V and LV ChP, with small numbers in the septum pellucidum and paraventricular zone 24 hrs following LPS ICV. Scale = 1mm.

**Supplemental figure 4.**
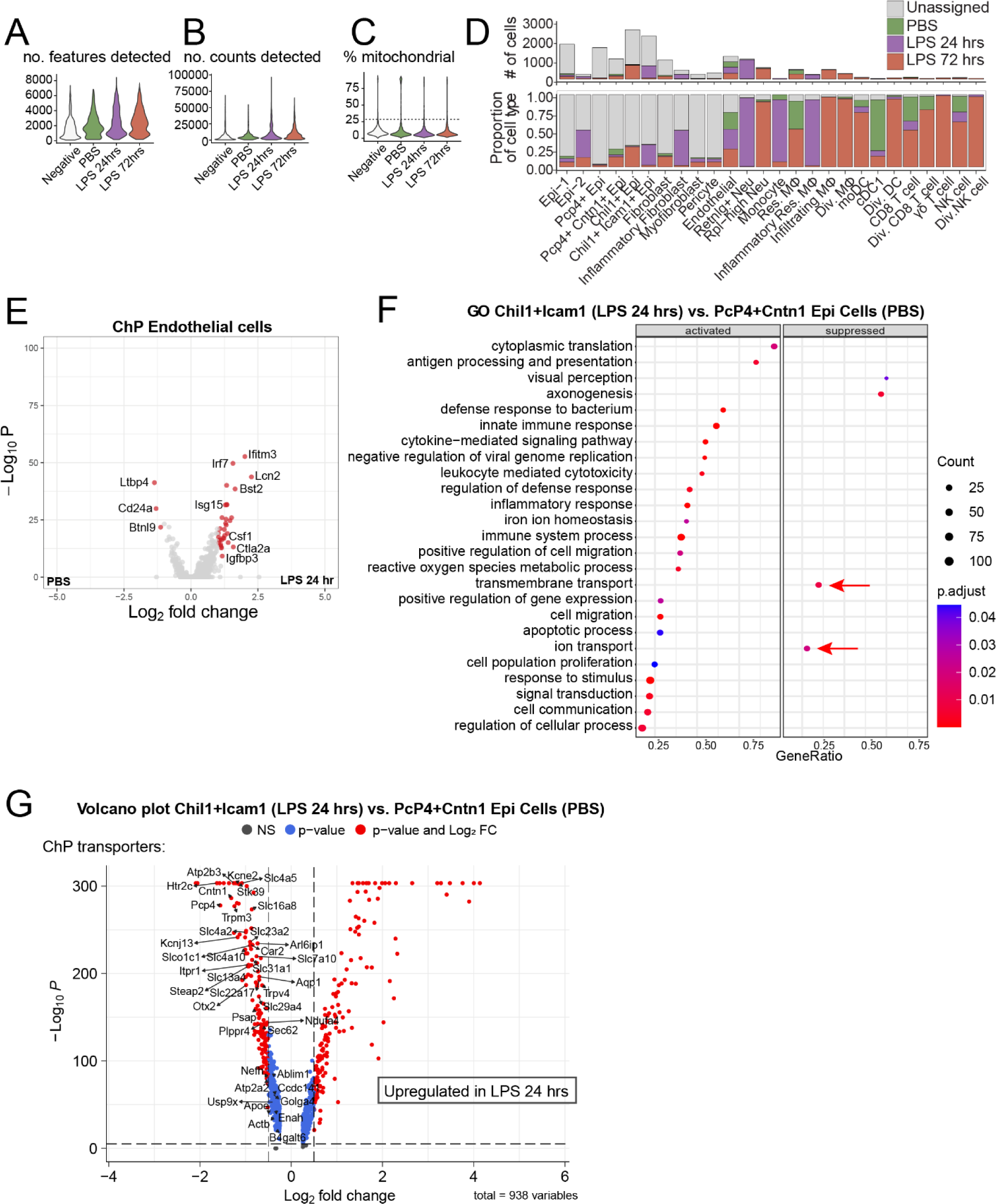
Supportive data showing neutrophil and monocytes infiltration through the ChP epithelial barrier. (**A**) Lyz2^ZsGreen^ mice had green fluorescence in S100A9+ neutrophils which were abundant in ChP explants 24 hours after LPS ICV. Scale = 100 µm. (**B**) Schematics and snapshots of two-photon in vivo imaging showing peripherally labeled Ly6G+ neutrophils present and moving in the ChP. Also see Supplemental video 3. Scale = 50 µm. (**C**) Representative electron microscopy images showing neutrophils attached to the blood vessel (top), within the ChP stroma (middle) and on the apical sides of ChP epithelial cells (bottom), 24 hours after LPS ICV. Scale = 2µm. (**D**) Examples of neutrophils in close contact with epithelial cells observed by ImageStream; (**E**) Representative gating image demonstrating neutrophil-epithelial population.

**Supplemental figure 5.**
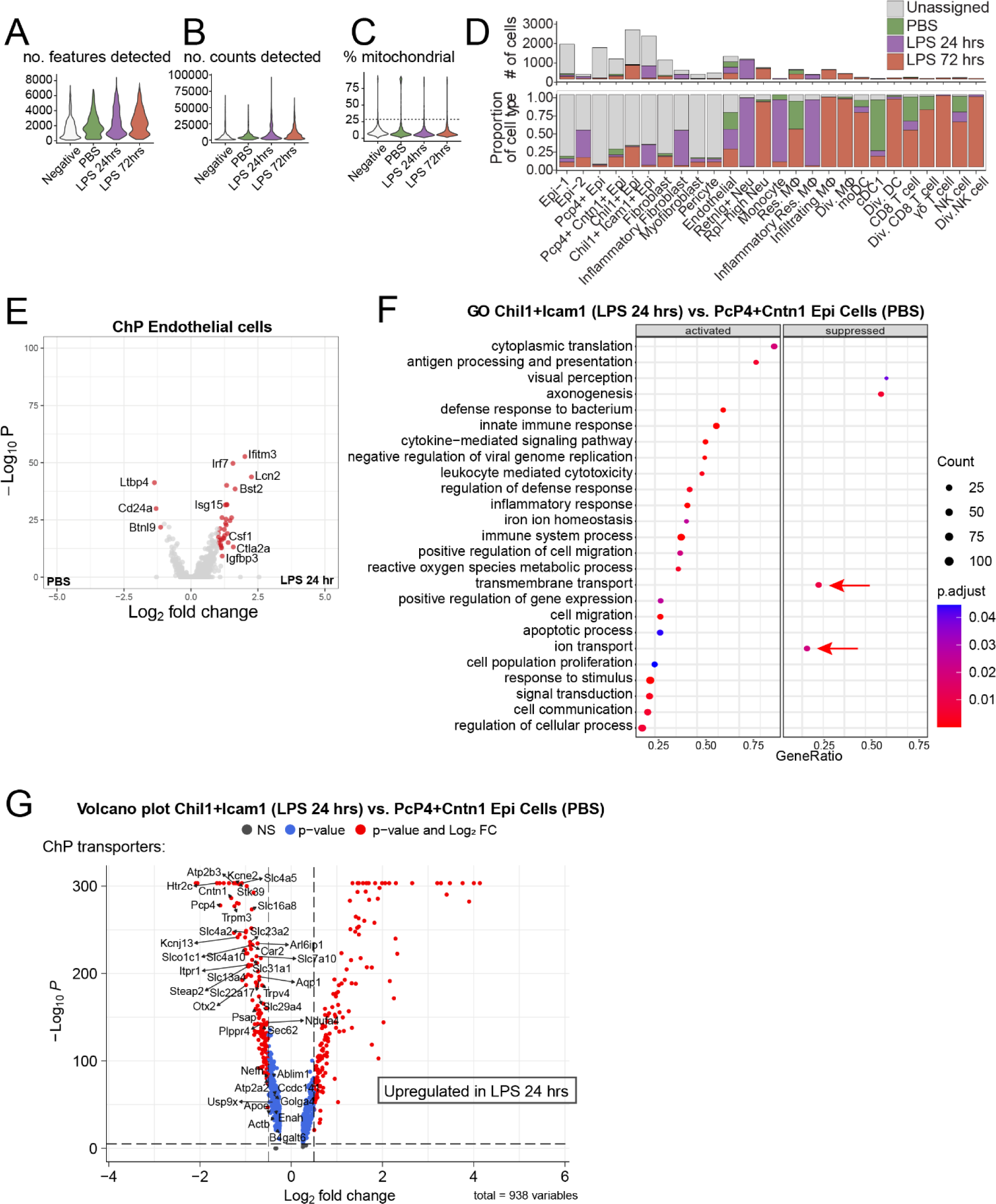
ChP scRNAseq data QC and differential gene expression analysis. (**A-C**) QC plots from ChP scRNAseq showing comparable gene detection, cell count, and cell viability across samples. (**D**) Cell frequencies and relative proportions by assigned identity. (**E**) Volcano plots showing differentially expressed genes between endothelial cells from PBS vs. LPS 24 hours; (**F**) GO-terms of differentially expressed genes between PBS PcP4+ Cntn1+ and 24 hours LPS Icam1+ Chil1+ epithelial cells, showing increased immune-related terms and decreased ion transport terms; (**G**) Volcano plot showing differentially expressed genes related to ion transport.

**Supplemental figure 6.**
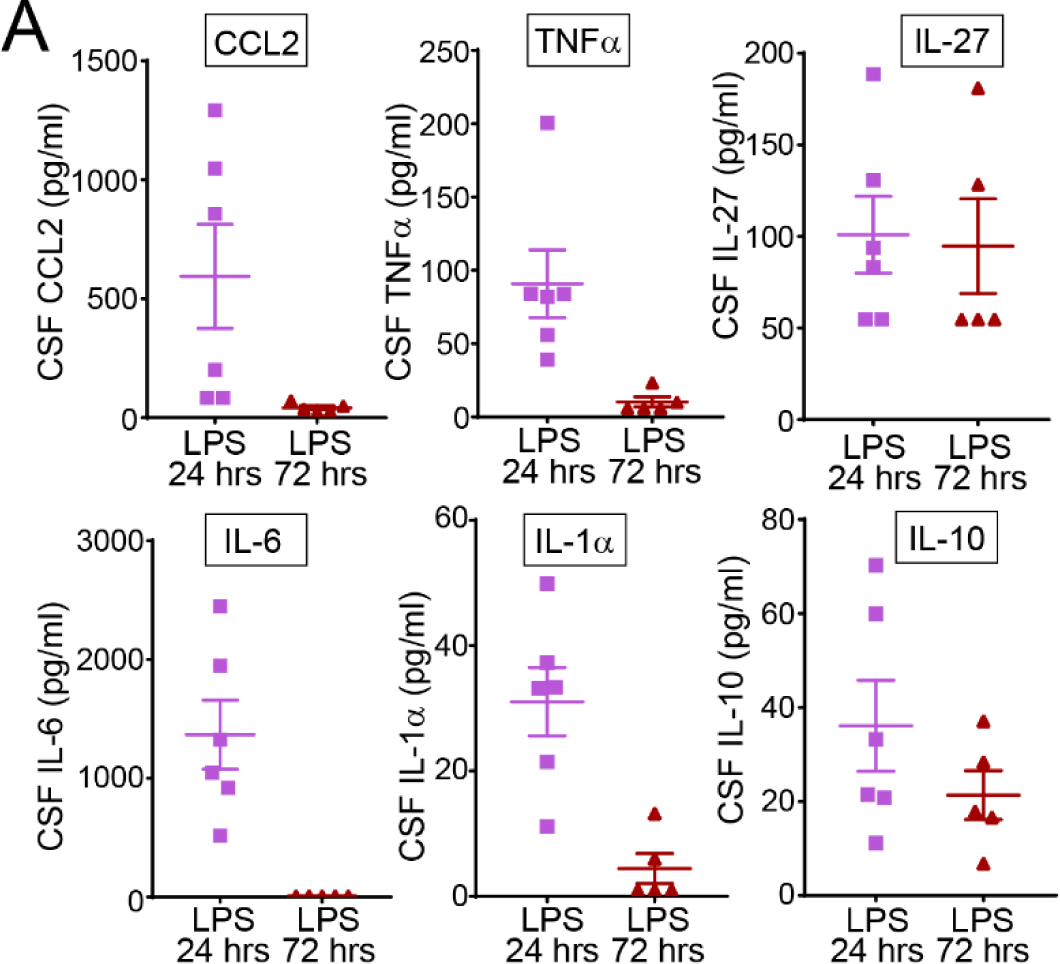
CSF cytokines and chemokines levels following LPS ICV by ELISA.

**Supplemental figure 7.**
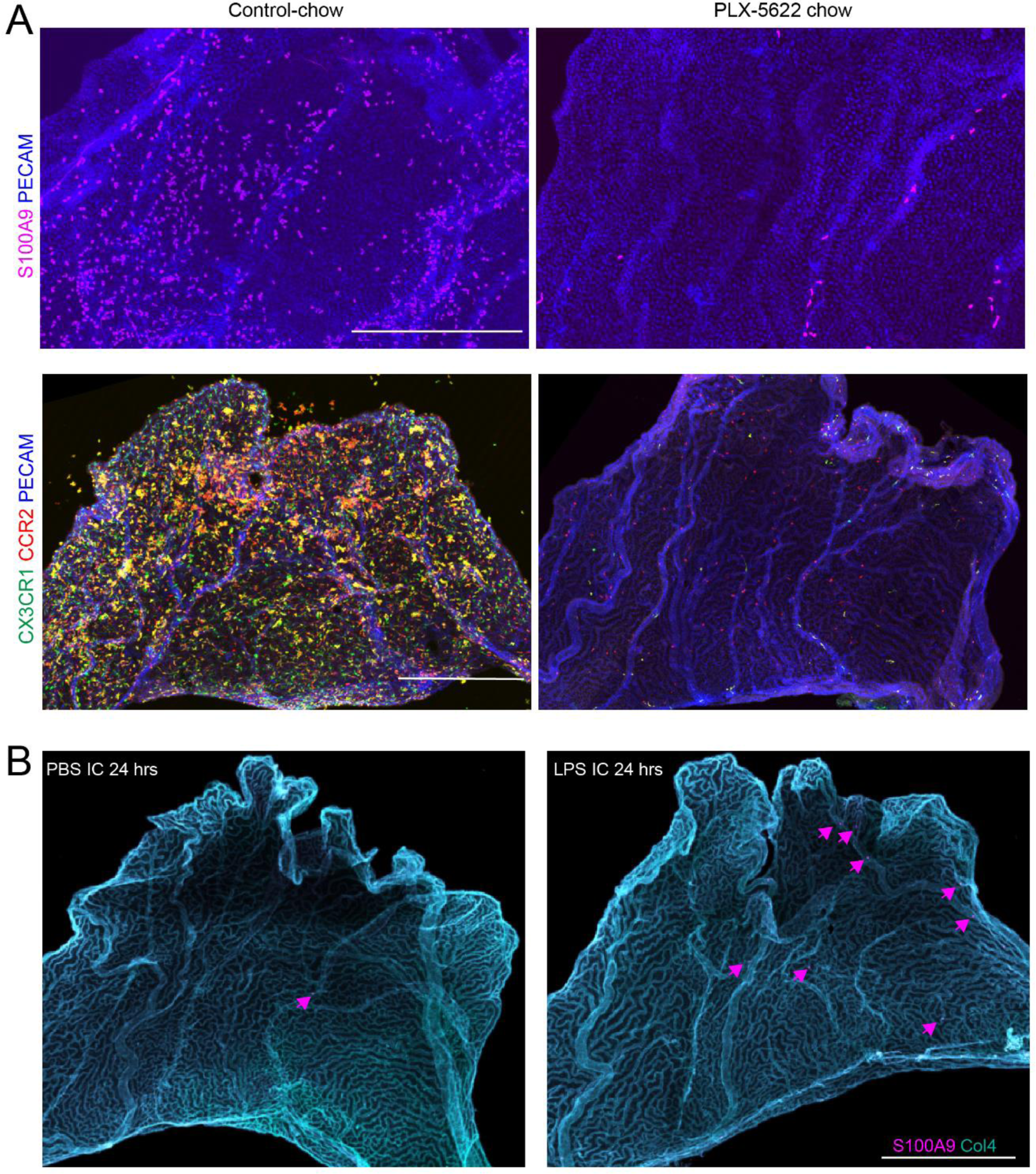
Direct ChP exposure to LPS is essential to initiate immune infiltration. (**A**) representative images showing mice with PLX-5622 chow diet failed to recruit neutrophils and monocytes following LPS ICV. Scale = 200 µm. (**B**) Representative images showing minimal S100A9+ leukocytes infiltration (purple arrows) into the ChP at 24 hours when LPS was delivered to the blood at the same dose as ICV. Scale = 500 µm.

**Supplemental figure 8.**
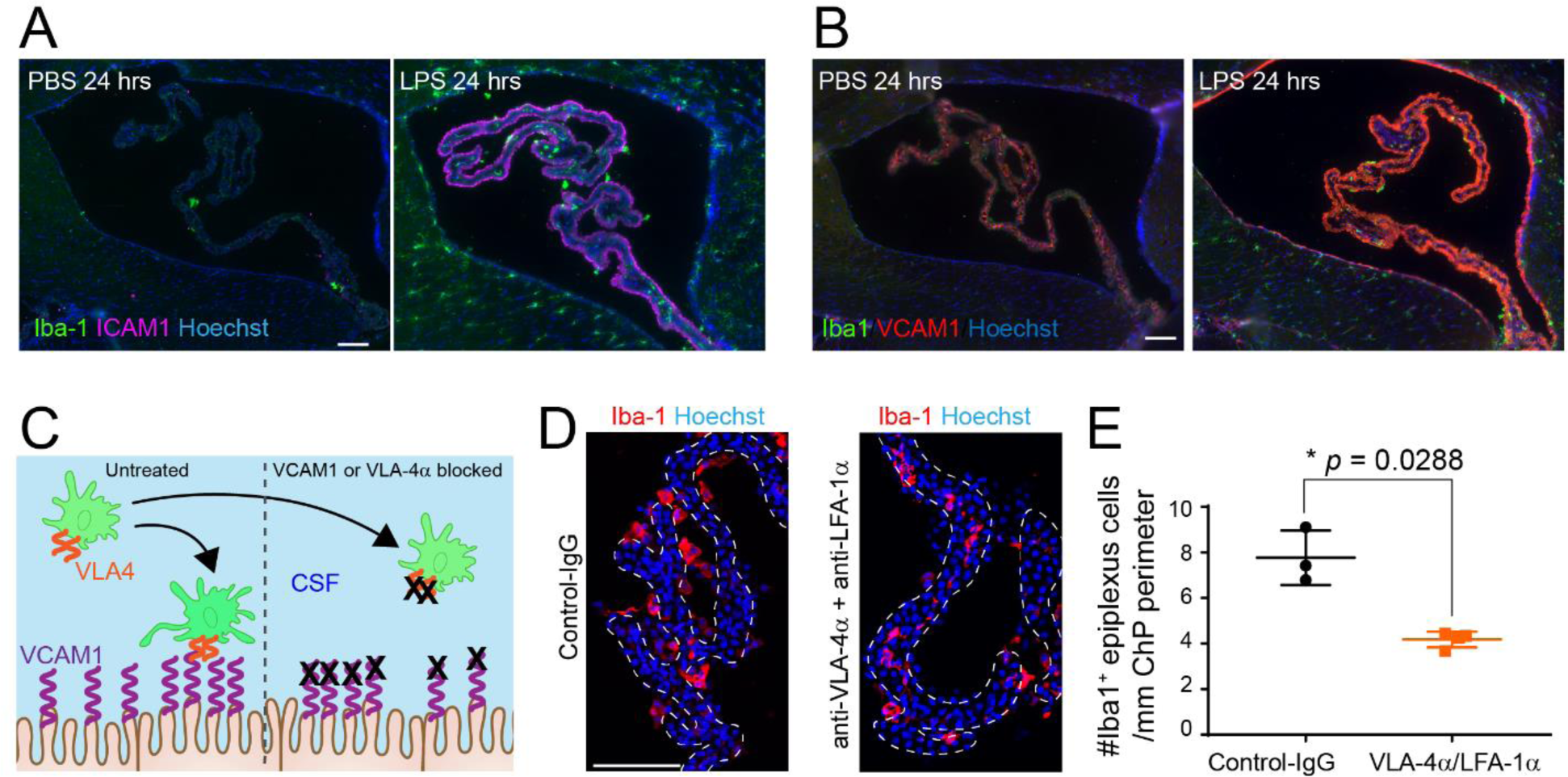
ChP epithelial adhesion molecules contribute to recruitment of epiplexus macrophages. (**A-B**) Representative images showing increased VCAM1 and ICAM1 expression on the ChP apical surface and ventricle wall 24 hours after LPS ICV. Scale = 100 µm. (**C-E**) Schematics and representative images and quantifications showing reduced numbers of total Iba1+ epiplexus macrophages in mice treated with anti-VLA-4 α /LFA-1α antibodies following LPS ICV (* *p* = 0.0288). Scale = 100 µm. Data are presented as mean ± SD.

**Supplemental figure 9.**
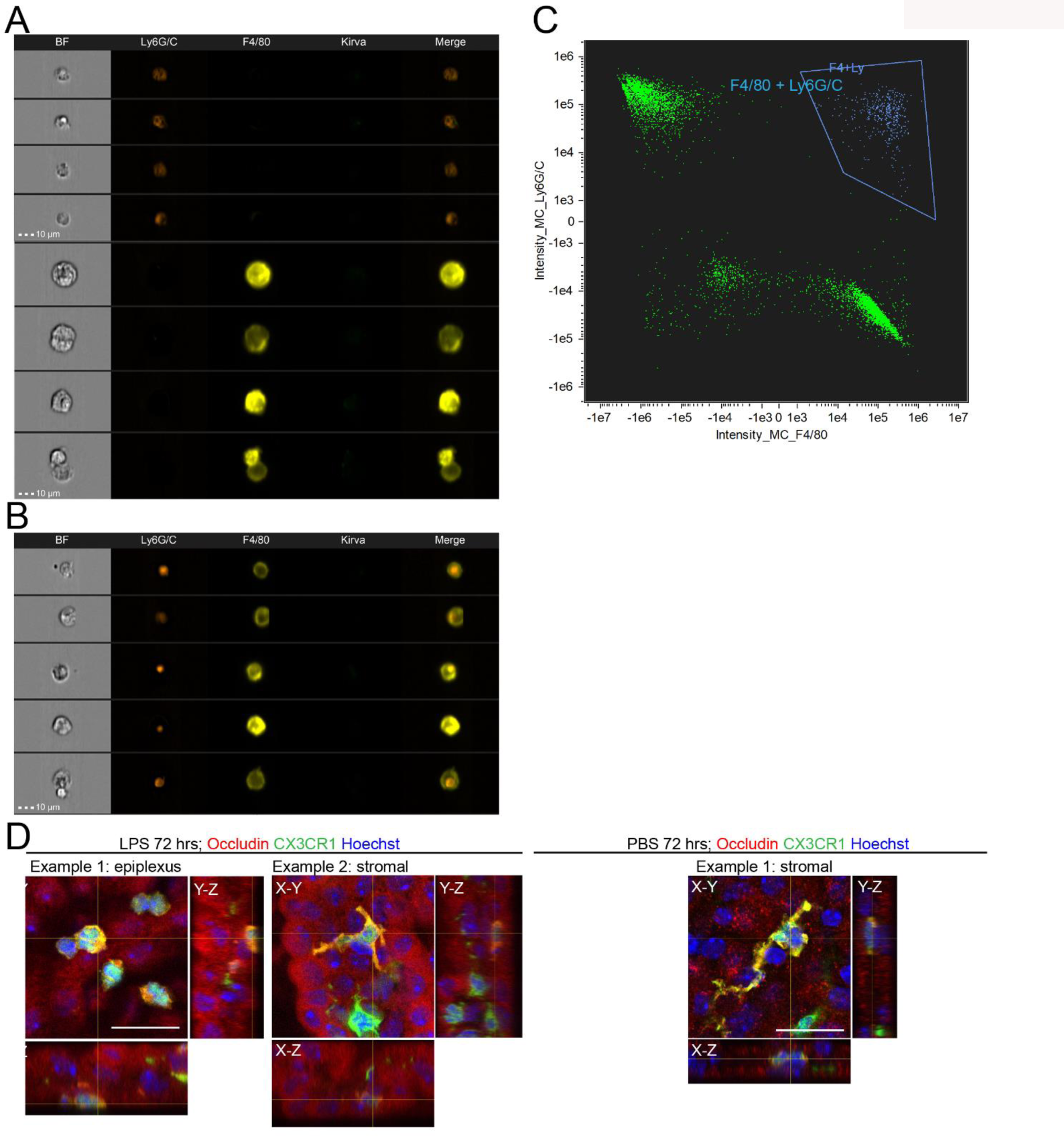
(**A**) Example cell images showing distinct staining and identification of neutrophils (Ly6G/C) and macrophages (F4/80) in the ChP by ImageStream. (**B**) Examples of neutrophils engulfed by macrophages. (**C**) Representative gating image demonstrating the cell population representing macrophages that engulfed neutrophils. (**D**) Additional representative 3D images and orthogonal views showing epiplexus and stromal CX3CR1+ macrophages that contain occludin in ChP explants from mice 72 hours after LPS ICV and. In PBS controls, occludin+ CX3CR1+ cells are predominantly stromal. Scale = 20 µm.

## Supplemental videos

**Video S1**. Timecourse montage showing LysM-ZsGreen neutrophils (green) entering the ChP over the course of 24 hours. Red: dextran-Texas Red. Scale = 100 µm.

**Video S2**. Representative video showing LysM-ZsGreen neutrophils (green) travel rapidly across CSF and on the ChP 24 hours following LPS ICV. Red: dextran-Texas Red. Scale = 100 µm.

**Video S3**. Representative video showing neutrophils peripherally labeled by IC Ly6G-568 antibody in the ChP. Red: Ly6G-568 (neutrophils); Green: Dextran-FITC. Scale = 50 µm.

**Video S4**. Representative montage showing 2 examples of increased number and mobility of CX3CR1^GFP^ myeloid cells (green) in the ChP 72 hours following LPS ICV. Red: dextran-Texas Red. Scale = 100 µm.

**Video S5**. Representative montage showing CX3CR1^GFP^ myeloid cells (green) traveling in the CSF 48 hours following LPS ICV (right), in comparison to baseline (left). Red: dextran-Texas Red. Scale = 100 µm.

**Video S6**. Montage showing 2 examples of CX3CR1^GFP^ myeloid cells (green) traveling from and towards the ventricle wall across multiple z-plains within one imaging stack of Supplementary Video V5. Red: dextran-Texas Red. Scale = 100 µm.

**Video S7**. Montage showing 3 examples of CX3CR1^GFP^ myeloid cells (green) traveling towards the ChP across multiple z-plains within one imaging stack of Supplementary Video V8. Red: dextran-Texas Red. Scale = 100 µm.

**Video S8**. Representative video showing one CX3CR1^GFP^ myeloid cells (green) traveling through CSF and landing on the ChP. Red: dextran-Texas Red. Scale = 100 µm.

**Video S9**. Representative video showing one CX3CR1^GFP^ myeloid cell (green) with large vacuoles inside its cytosol. Video was taken 24 hours following LPS ICV. Scale = 25 µm.

**Video S10**. Representative video showing CX3CR1^GFP^ myeloid cells (green) converging into one large clump. Video was taken 72 hours following LPS ICV. Scale = 25 µm.

**Video S11**. Representative video showing epiplexus CX3CR1^creER-TdTomato^ myeloid cells (red) moving on the apical surface of the ChP, judging based on TTR^mNeon^ labeling of epithelial cells. Video was taken 48 hours following LPS ICV. Scale = 100 µm.

## Supplemental datasets

**Dataset S1.** CSF scRNAseq celltype and subtype markers.

**Dataset S2.** ChP scRNAseq celltype and subtype markers.

