## Supplementary material for "A collaboration between immune cells and the choroid plexus epithelium in brain inflammation": MATLAB code: ChP_red_green_Colocalization.m

```
javaaddpath 'C:\Program Files\MATLAB\R2018a\java\mij.jar'
javaaddpath 'C:\Program Files\MATLAB\R2018a\java\ij-1.52a.jar'
javaaddpath 'C:\Users\LehtinenLab\Dropbox\AndermannLab\users\Fred\TurboRegHL_.jar'
 
fdir = 'G:\Michaela\HuixinJinn\Colocalization\For_Lab_Meeting';

greenname = 'LPS_3_day_GFP';
redname = 'LPS_3_day_RFP';

greenmov = pipe.io.readTiff(strcat(fdir,filesep,greenname,'.tif'));
redmov = pipe.io.readTiff(strcat(fdir,filesep,redname,'.tif'));
Nx = size(greenmov,1);
Ny = size(greenmov,2);
Nz = size(greenmov,3);
%%
mova(1,:,:,:) = redmov;
mova(2, :, :, :) = greenmov;
implay2chan(mova);
%%
movb = rescale(redmov) .* rescale(greenmov);
implay(rescale(movb));
%%
testyellow = imbinarize(movb);
%%
redmov2 = rescale(redmov);
redslice = redmov2(: ,:,9);
red_low = prctile(redslice(:),70);
red_hi = prctile(redslice(:),99);
figure
imshow(rescale(redslice));
figure
imshow(rescale(imadjust(redslice,[red_low,red_hi])));
%%
bintest = imbinarize(rescale(imadjust(redslice,[red_low,red_hi])));
%%
pctlemovg = zeros(Nx, Ny, Nz);
pctlemovr = zeros(Nx, Ny, Nz);
redmov2 = rescale(redmov);
greenmov2 = rescale(greenmov);
for i=1:Nz
    redslice = redmov2(: ,:,i);
    red_low = prctile(rescale(redmov(:)),75);
    red_hi = prctile(rescale(redmov(:)),100);
    greenslice = greenmov2(: ,:,i);
    green_low = prctile(rescale(greenmov(:)),75);
    green_hi = prctile(rescale(greenmov(:)),100);
    greenslice = imadjust(greenslice,[green_low,green_hi]);
    redslice = imadjust(redslice,[red_low,red_hi]);
    pctlemovg(:, :, i) = greenslice;
    pctlemovr(:, :, i) = redslice;   
end

BWG = imbinarize(pctlemovg);
BWR = imbinarize(pctlemovr);
%%
video = zeros(Nx, Ny, 3, Nz);
video2 = zeros(Nx, Ny, 3, Nz);
green_tot = 0;
red_tot = 0;
yellow_tot = 0;
for i=1:Nz
    bwfiltg = bwareafilt(BWG(:, :, i), [200,inf]);
    bwfiltr = bwareafilt(BWR(:, :, i), [200,inf]);
    green = bwarea(bwfiltg);
    red = bwarea(bwfiltr);
    
    green_tot = green_tot + green;
    red_tot = red_tot + red;
    
    coloc = bitand(bwfiltg, bwfiltr);
    coloc = bwareafilt(coloc, [200, inf]);
    grn = bitxor(bwfiltg, coloc);
    rd = bitxor(bwfiltr, coloc);
    
    yellow = bwarea(coloc);
    yellow_tot = yellow_tot + yellow;
    
    rgbImage = cat(3, bwfiltr , bwfiltg , video(:,:,1,i));
    videoy(:, :, :, i) = rgbImage;
    
    rgbImage = cat(3, rd , grn , coloc);
    videob(:, :, :, i) = rgbImage;
end
yellow_percent = yellow_tot/(green_tot + red_tot - yellow_tot);
%%
max_projg = max(pctlemovg, [], 3);
max_projr = max(pctlemovr, [], 3);
max_bwg = imbinarize(max_projg);
max_bwr = imbinarize(max_projr);
max_bwg = bwareafilt(max_bwg, [200,inf]);
max_bwr = bwareafilt(max_bwr, [200,inf]);
max_bwy = bitand(max_bwg, max_bwr);
max_bwy = bwareafilt(max_bwy, [200,inf]);
new_green = bitxor(max_bwg, max_bwy);
new_green = bwareafilt(new_green, [200,inf]);
new_red = bitxor(max_bwr, max_bwy);
new_red = bwareafilt(new_red, [200,inf]);
labelg = bwlabel(new_green);
labelr = bwlabel(new_red);
labely = bwlabel(max_bwy);
ngreen = max(labelg(:));
nred = max(labelr(:));
nyellow = max(labely(:));
%% Image Processing
fiberboig = fibermetric(uint16(greenmov));
fiberboir = fibermetric(uint16(redmov));
blurg = imgaussfilt(greenmov, 5);
blurr = imgaussfilt(redmov, 5);
%% Blending
newmovg = zeros(Nx, Ny, Nz);
newmovr = zeros(Nx, Ny, Nz);
for i=1:Nz
  fiberboi2g = imadd(double(blurg(:,:,i)), double(fiberboig(:, :, i)));
  fiberboi2r = imadd(double(blurr(:,:,i)), double(fiberboir(:, :, i)));  
  newmovg(:, :, i) = fiberboi2g;
  newmovr(:, :, i) = fiberboi2r;
end
%%
newmovg = rescale(newmovg);
newmovr = rescale(newmovr);
gmid = max(newmovg(:, :, Nz/2-1:Nz/2+1), [], 3);
rmid = max(newmovr(:, :, Nz/2-1:Nz/2+1), [], 3);
greenthresh = graythresh(gmid);
redthresh = graythresh(rmid);
BWGtest = imbinarize(gmid, greenthresh);
BWG = imbinarize(rescale(newmovg), greenthresh);
BWR = imbinarize(rescale(newmovr), redthresh);
%%
video = zeros(Nx, Ny, 3, Nz);
green_tot = 0;
red_tot = 0;
yellow_tot = 0;
for i=1:Nz
    bwfiltg = bwareafilt(BWG(:, :, i), [100,inf]);
    bwfiltr = bwareafilt(BWR(:, :, i), [100,inf]);
    green = bwarea(bwfiltg);
    red = bwarea(bwfiltr);
    
    green_tot = green_tot + green;
    red_tot = red_tot + red;
    
    coloc = bitand(bwfiltg, bwfiltr);
    grn = bitxor(bwfiltg, coloc);
    rd = bitxor(bwfiltr, coloc);
    
    yellow = bwarea(coloc);
    yellow_tot = yellow_tot + yellow;
    
    rgbImage = cat(3, rd , grn , coloc);
    video(:, :, :, i) = rgbImage;
end
yellow_percent = yellow_tot/(green_tot + red_tot - yellow_tot);
%%
max_projg = max(newmovg, [], 3);
max_projr = max(newmovr, [], 3);
max_bwg = imbinarize(max_projg);
max_bwr = imbinarize(max_projr);
max_bwg = bwareafilt(max_bwg, [100,inf]);
max_bwr = bwareafilt(max_bwr, [100,inf]);
max_bwy = bitand(max_bwg, max_bwr);
max_bwy = bwareafilt(max_bwy, [100,inf]);
new_green = bitxor(max_bwg, max_bwy);
new_green = bwareafilt(new_green, [100,inf]);
new_red = bitxor(max_bwr, max_bwy);
new_red = bwareafilt(new_red, [100,inf]);
labelg = bwlabel(new_green);
labelr = bwlabel(new_red);
labely = bwlabel(max_bwy);
ngreen = max(labelg(:));
nred = max(labelr(:));
nyellow = max(labely(:));
%%
new_mov_g = zeros(150, 150, ngreen);
new_mov_bin_g = zeros(150, 150, ngreen);
for i=1:ngreen
    segim = zeros(Nx, Ny);
    segim(labelg == i) = 1;
    s = regionprops(segim, 'Centroid');
    xLeft = s.Centroid(1) - 75;
    width = 149;
    yTop = s.Centroid(2) - 75;
    height = 149;
    croppedImage = imcrop(max_projg, [xLeft, yTop,width, height]);
    croppedBin = imcrop(segim, [xLeft, yTop,width, height]);
    if size(croppedImage) == size(new_mov_g(:, :, 1))
        new_mov_g(:, :, i) = croppedImage;
        new_mov_bin_g(:, :, i) = croppedBin;
    end
end

%% Test segmentations
new_mov = zeros(150, 150, length(new_X));
new_mov_seg = zeros(150, 150, length(new_X));
new_mov_bin = zeros(150, 150, length(new_X));
for i=1:length(new_X)
    xLeft = new_X(i) - 75;
    width = 149;
    %width = xLeft + 2 * 50;
    yTop = new_Y(i) - 75;
    height = 149;
    %height = yTop + 2 * 50;
    croppedImage = imcrop(mov, [xLeft, yTop,width, height]);
    croppedSeg = imcrop(I1, [xLeft, yTop,width, height]);
    croppedBin = imcrop(segments, [xLeft, yTop,width, height]);
    if size(croppedImage) == size(new_mov(:, :, 1))
        new_mov(:, :, i) = croppedImage;
        new_mov_seg(:, :, i) = croppedSeg;
        new_mov_bin(:, :, i) = croppedBin;
    end
end
```
