## Supplementary material for "A collaboration between immune cells and the choroid plexus epithelium in brain inflammation": MATLAB code: PostHocMatching.m

```
fdir = '\\169.254.231.223\DATA\Huixin and Jin\Sample images for Fred and Michaela\Two-P and explant view matching test';
fname_2p = 'C1-0118-0425-017-baseline_10x'; %2p name
fname_ex = 'C2-LysM-ZsGreen post-LPS 017'; %.tif name %HX .czi name? Match with line 50. czi or tif
%% shape and scale 2p image (ASSUMING .TIF STACK WITH METADATA)
im_2p = pipe.io.read_tiff(strcat(fdir,filesep,fname_2p,'.tif'));
im_2p_info = imfinfo(strcat(fdir,filesep,fname_2p,'.tif'));
im_info = im_2p_info(1);
res_2p = im_info(1).XResolution;

%REMOVE THIS LATER; ONLY MEANT TO CORRECT MIS-LABELED OBJECTIVE ZOOM
% res_2p = res_2p * (10/15);
%

%parse extra metadata
temp = split(im_info(1).ImageDescription,[newline,"="]);
temp = reshape(temp(1:end-1),2,[]);
map_2p = containers.Map(temp(1,:),temp(2,:));

Nx_2p = im_info(1).Width;
Ny_2p = im_info(1).Height;
Nz_2p = str2num(map_2p('slices'));
Nt_2p = str2num(map_2p('frames'));

%reshape and scale based on metadata
im_2p = reshape(im_2p,Nx_2p,Ny_2p,Nz_2p,Nt_2p);
im_2p_proj = mean(im_2p,4);
im_2p_proj = max(im_2p_proj,[],3);
im_2p_proj = imresize(im_2p_proj,1/res_2p);
im_2p_proj = rescale(im_2p_proj);
%% load, shape, and scale explant image, .tif stack
% im_ex = pipe.io.read_tiff(strcat(fdir,filesep,'test3.tif'));
% im_ex_info = imfinfo(strcat(fdir,filesep,'test3.tif'));
% im_ex_info = im_ex_info(1);
% res_ex = im_ex_info(1).XResolution;
% 
% temp = split(im_ex_info(1).ImageDescription,[newline,"="]);
% temp = reshape(temp(1:end-1),2,[]);
% map_ex = containers.Map(temp(1,:),temp(2,:));
% 
% Nx_ex = im_ex_info(1).Width;
% Ny_ex = im_ex_info(1).Height;
% Nz_ex = str2num(map_ex('slices'));
% 
% % im_ex = reshape(im_ex,Nx_ex,Ny_ex,Nz_ex);
% im_ex_proj = max(im_ex,[],3);
% im_ex_proj = imresize(im_ex_proj,1/res_ex);
% im_ex_proj = rescale(im_ex_proj);

% %% load, shape, and scale explant image (ASSUMING .CZI WITH METADATA)
reader = bfGetReader(strcat(fdir,filesep,fname_ex,'.tif'));
M = reader.getMetadataStore();
Nc_ex = M.getPixelsSizeC(0).getValue();
Nx_ex = M.getPixelsSizeX(0).getValue();
Ny_ex = M.getPixelsSizeY(0).getValue();
Nz_ex = M.getPixelsSizeZ(0).getValue();
res_ex = 1 / double(M.getPixelsPhysicalSizeX(0).value);

cr = 1:Nc_ex;
zr = 1:Nz_ex;

im_exp = zeros(numel(cr),Ny_ex,Nx_ex,numel(zr),'uint16');

for c = cr
    for z = zr
        idx = reader.getIndex(z-1,c-1,0)+1;
        im_exp(c,:,:,z) = bfGetPlane(reader,idx);
    end
end

im_ex_proj = squeeze(max(im_exp(1,:,:,:),[],4));
im_ex_proj = imresize(im_ex_proj,1/res_ex); %make it unit scale (i.e. 1um = 1px)
im_ex_proj = rescale(im_ex_proj); %rescale the brightness values
%% use GUI to do rough alignment
matchapp = ManualCompareImages(im_2p_proj, im_ex_proj);
waitfor(matchapp,'closeflag');
flip = matchapp.imflip;
theta = matchapp.theta;
delete(matchapp);

%performs necessary flip and rotation
im_ex_proj = imrotate(im_ex_proj,theta,'crop');
if flip
    im_ex_proj = fliplr(im_ex_proj);
end
%% select points from both images using point select GUI
[movingPoints, fixedPoints] = cpselect(im_2p_proj,im_ex_proj,'Wait',true);
tform = fitgeotrans(movingPoints, fixedPoints,'projective');

%% make binary image of 2p to mask 
I = imgaussfilt(im_2p_proj,20);
I_bin = I > mean(I(:));
I_bin = imopen(I_bin, strel('disk',30));
I_bin = imfill(I_bin,'holes');

%% fig transformation
im_2p_warp = imwarp(im_2p_proj,tform,'OutputView', imref2d(size(im_ex_proj)));
I_bin_warp = imwarp(I_bin,tform,'OutputView', imref2d(size(im_ex_proj)));

im_2p_warp = imgaussfilt(im_2p_warp,3);
im_ex_proj = imgaussfilt(im_ex_proj,3);

im_2p_warp_masked = imadjust(im_2p_warp.*I_bin_warp);
im_ex_proj_masked = imadjust(im_ex_proj.*I_bin_warp);

%% do demons and show match
[~,im_2p_demons] = imregdemons(im_2p_warp_masked,im_ex_proj_masked,'AccumulatedFieldSmoothing',2.5,'PyramidLevels',9);
figure,imshow(im_2p_demons);
figure,imshowpair(im_2p_demons,im_ex_proj_masked);

%% show matching points and overlay
figure;
subplot(1,5,1:3);
showMatchedFeatures(imadjust(im_2p_proj), im_ex_proj, movingPoints, fixedPoints,'montage','PlotOptions', {'rx', 'rx', 'y'});
subplot(1,5,4:5);
imshowpair(im_2p_demons,im_ex_proj_masked);
```
